# NRCAM variant defined by microexon skipping is a targetable cell surface proteoform in high-grade gliomas

**DOI:** 10.1101/2025.01.09.631916

**Authors:** Priyanka Sehgal, Ammar S. Naqvi, Makenna Higgins, Jiageng Liu, Kyra Harvey, Julien Jarroux, Taewoo Kim, Berk Mankaliye, Pamela Mishra, Grace Watterson, Justyn Fine, Jacinta Davis, Katharina E. Hayer, Annette Castro, Adanna Mogbo, Charles Drummer, Daniel Martinez, Mateusz P. Koptyra, Zhiwei Ang, Kai Wang, Alvin Farrel, Mathieu Quesnel-Vallieres, Yoseph Barash, Jamie B. Spangler, Jo Lynne Rokita, Adam C. Resnick, Hagen U. Tilgner, Thomas De Raedt, Daniel J Powell, Andrei Thomas-Tikhonenko

**Author notes:** Département d’immunologie et de biologie cellulaire, Université de Sherbrooke, Sherbrooke, QC. Center for Cancer and Immunology Research, Children’s National Hospital, Washington, DC. Senior author (Andrei Thomas-Tikhonenko). Twitter: @andrei_thomas_t (Andrei Thomas-Tikhonenko).

## Abstract

To overcome the paucity of known tumor-specific surface antigens in pediatric high-grade glioma (pHGG), we contrasted splicing patterns in pHGGs and normal brain samples. Among alternative splicing events affecting extracellular protein domains, the most pervasive alteration was the skipping of ≤30 nucleotide-long exons. Several of these skipped microexons mapped to L1-IgCAM family members, such as *NRCAM*. Bulk and single-nuclei short- and long-read RNA-seq revealed uniform skipping of *NRCAM* microexons 5 and 19 in virtually every pHGG sample. Importantly, the Δex5Δex19 (but not the full-length) NRCAM proteoform was essential for pHGG cell migration and invasion in vitro and tumor growth in vivo. We developed a monoclonal antibody selective for Δex5Δex19 NRCAM and demonstrated that “painting” of pHGG cells with this antibody enables killing by T cells armed with an FcRI-based universal immune receptor. Thus, pHGG-specific NRCAM and possibly other L1-IgCAM proteoforms are promising and highly selective targets for adoptive immunotherapies.

## INTRODUCTION

Pediatric high-grade gliomas (pHGG) are some of the most recalcitrant, chemoresistant, and often surgically unresectable childhood cancers. For example, children affected by diffuse midline gliomas (DMG) nearly universally succumb to the disease within 8-10 months of diagnosis ^1^. Its adult counterpart, glioblastoma multiforme (GBM), is equally lethal ^2^. Currently, there are no effective standard of care therapies for these patients, despite decades of molecularly agnostic clinical trials, underscoring the importance of developing novel immunotherapies. Several targets for CAR T cells in GBM have been developed, most notably EGFRvIII ^3,4^, HER2 ^5^, IL13Rα2 ^6^, and Eph2A ^7^, but so far the corresponding immunotherapeutics enjoyed limited success in the clinic, owing in part to the frequent emergence of antigen escape variants [reviewed in ^8–11^].

Although GBMs and pHGGs are thought to arise in the same lineage, the latter express somewhat different repertoires of surface antigens. Validated pHGG targets include GD2 and B7-H3 [reviewed in ^12,13^]. GD2 in particular is highly expressed by H3K27M-mutated DMG cells ^14^, and the first 4 DMG patients were recently treated with anti-GD2 CAR, with some clinical and radiographic improvement ^15^. Similarly, both anaplastic astrocytomas and DMG express B7-H3 (a.k.a. CD276), and an anti-B7-H3 CAR has shown activity in preclinical models of pediatric tumors ^16^. However, targeting GD2 is associated with well-documented (although not universally observed) on-target, off-tumor toxicities ^17,18^, and side effects of another anti-B7-H3 CAR have been only assessed in the mouse ^19^ and in a relatively small cohort of human patients with DIPG ^20,21^.

Even if GD2- and/or B7-H3-directed immunotherapies become standards of care, the clinical use of CAR T cells in patients with hematologic malignancies suggests that there is no such thing as the “perfect” target, and there cannot be too many alternatives ^22,23^, the more cancer-specific the better ^24^. Here we describe our effort to systemically identify new targets using the pHGG model, where mutation burden is typically low, and aberrant mRNA splicing is a key post-transcriptional mechanism of proteome diversification ^25^. This effort has led to the identification of the four members of the L1-immunoglobulin (Ig) sub-family of cell adhesion molecules (CAMs), in particular NRCAM splice variant, as promising targets for adoptive T cell therapy.

## RESULTS

### Identification of microexon-derived surface proteoforms in pHGG

Our initial analysis of 325 pHGG tumors from the Open Pediatric Brain Tumor Atlas ^26^ revealed that not only is aberrant splicing prevalent in pHGG, but these tumors had some of the most heterogeneous splicing programs ^25^. Thus, the rMATS-turbo splicing algorithm ^27^ was used to compare 184 pHGG patient samples to each of the 7 healthy controls comprised of an adult brain homogenate, an adult brain stem, an adult cerebellum, an adult occipital cortex, two fetal brains, and one pediatric normal cortex. pHGG-specific splice junction candidates were filtered for single-exon (SE) events, whose junctions were mapped to predicted protein structures using UniProt annotations and filtered for SE events corresponding to extracellular protein domains (Figure 1A), to enable future development of therapeutic antibodies and chimeric antigen receptors.

**Figure 1.**
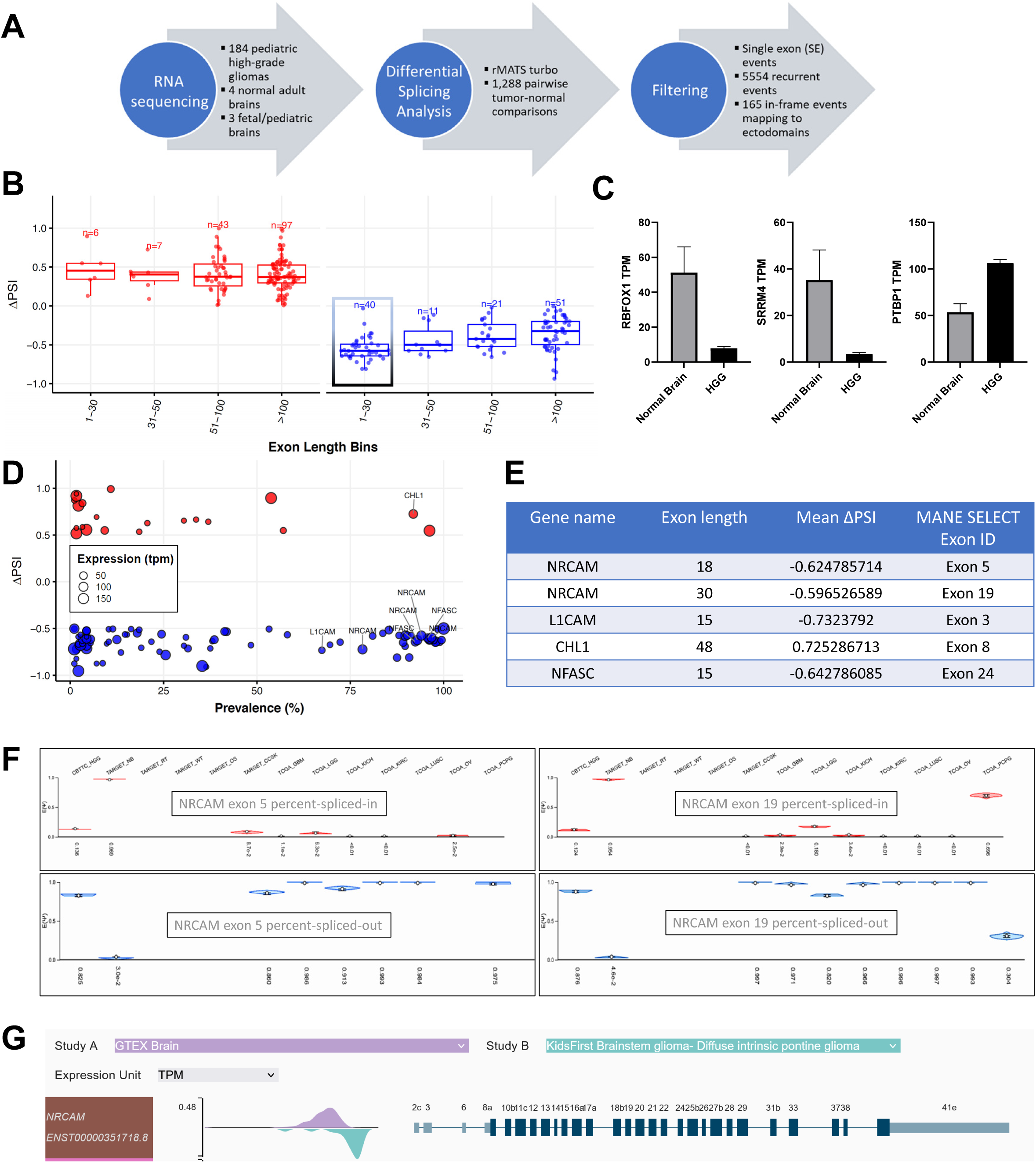
Aberrantly spliced surface protein-encoding transcripts in pHGG. **A.** The pipeline used to identify pHGG-specific splicing events. **B.** Difference in exon inclusion level (ΔPSI) between pHGG and normal brain samples (y axis). Splicing events were binned based on exon lengths (x axis). Small box indicates the only bin with >50% difference in absolute PSI values. **C.** Bar graphs showing the expression of select splice factors in normal brain vs. pHGG tissues. Each bar represents mean ± SEM. **D.** Scatter plot showing ΔPSI values, event prevalence, and transcript levels (TPM) of microexons differentially spliced in pHGG. **E.** Top aberrantly spliced pHGG microexons. **F.** Graphical output of MAJIQlopedia showing average PSIs of NRCAM exons 5 and 19 across tumor types represented in TCGA and CBTTC repositories**. G.** Visualization of the ENST00000351718.8 transcript and its expression level in normal brain (purple) vs Kids First Brainstem glioma (teal) using Xena portal.

These events were filtered for ≥40% prevalence and further binned according to exon lengths. The only events with mean differences in percent-spliced-in (ΔPSI) absolute values greater than 0.50 were skipping of very short exons: ≤30- and 31-50-nucleotide long (Figure 1B, box). These are known as microexons ^28^, and their dysregulation has been previously implicated in several CNS diseases ^29^, including autism spectrum disorder ^30^. Inclusion of microexons requires dedicated RNA binding proteins such as RBFOX ^31,32^ and SRRM4 ^30,33–35^, whose activity is counteracted by PTBP1, a promoter of microexon skipping ^36–38^. Of note, expression levels of *RBFOX1* and *SRRM4* mRNA were lower, and that of *PTBP1* were higher in pHGG compared to normal brain tissues (Figure 1C), which could explain the observed bias toward microexon skipping. To validate these findings in patient-derived samples, we performed qRT-PCR assays on pooled normal brain RNAs vs. 6 cultured pHGG patient-derived xenografts (PDXs). We observed statistically significant differences in *RBFOX1, SRRM4*, and *PTBP1* levels, which mirrored those seen in the original datasets (Supplemental Figure 1A).

Thus, we re-analyzed the entire rMATS-turbo output focusing on microexons corresponding to cell surface proteins and with |ΔPSI|≥0.5. We found that mRNAs encoding the 4 members of the L1-immunoglobulin superfamily of cell adhesion molecules (IgCAM), *L1CAM* itself, *NFASC*, *CHL1*, and *NRCAM*, were among the most profoundly and consistently mis-spliced transcripts (Figure 1D,E). In fact, neuronal cell adhesion molecule (*NRCAM*) mRNA had two microexon-related events: skipping of the 18-nt microexon 5 and the 30-nt microexon 19 [MANE nomenclature ^39^; exons 9 and 23 in GTEx v8 ^40^] (Figure 1E). We also observed the skipping of microexon 7 in *L1CAM* and microexon 24 - in *NFASC*. One exception to the trend toward exon skipping was increased inclusion of exon 8 in *CHL1*, but that exon was slightly longer, 48 nucleotides, which is borderline for the purpose of microexon definition ^29^. To validate these findings in patient-derived samples, we performed semi-quantitative, low-cycle-number RT-PCR assays on the samples from panel S1A. Using forward and reverse primers corresponding to exons flanking microexons in question, we were able to detect both skipping and inclusion events in the same reactions. Without exemption, the splicing patterns of the 6 PDXs matched those predicted by rMATS-Turbo (Supplemental Figure 1B).

Focusing on *NRCAM* as the most profoundly affected transcript, we asked whether its microexons are skipped in other human tumors. We queried MAJIQlopedia, our recently developed compendium of splicing variations across 86 human tissues and 41 cancer datasets ^41^. We found that skipping of *NRCAM* exons 5 and 19 was prevalent in all tumor types with detectable *NRCAM* splice junctions. They included pHGG, adult low-grade gliomas and GBMs, pheochromocytomas and paragangliomas, and also cancers of non-neural origins, such lung adenocarcinomas (Figure 1F). The sole exception was pediatric neuroblastoma, where *NRCAM* exons 5 and 19 were uniformly included. Of note, these splicing quantifications were based on MAJIQ ^42^, an algorithm orthogonal to rMATS-turbo, increasing our confidence in *NRCAM*-centric events. Further evidence for preferential *NRCAM* exon 5 and 19 skipping in pHGG was found in the output of Toil, yet another orthogonal approach to transcript reconstitution from short-read RNA-seq data ^43^. Using Xena portal ^44^, we observed that transcript ENST00000351718.8 lacking exons 5 and 19 was overexpressed in brainstem gliomas compared to normal brain tissues (Figure 1G). ENST00000351718.8 was also upregulated in GBMs, consistent with the MAJIQlopedia output (Supplemental Figure S2A).

### HGG-specific expression of NRCAM ***Δ***ex5***Δ***ex19 isoform

To determine whether microexon skipping reflects, at least to some extent, pHGG lineage, we included in the analysis the RNA-seq dataset corresponding to 20 human primary astrocyte samples, 1 neuron sample, and 4 oligodendrocytes ^45^. We observed that skipping of HGG-specific microexons occurred primarily in astrocytes and oligodendrocytes, but not neurons, consistent with the pHGG eponymous cell of origin (Figure 2A).

**Figure 2.**
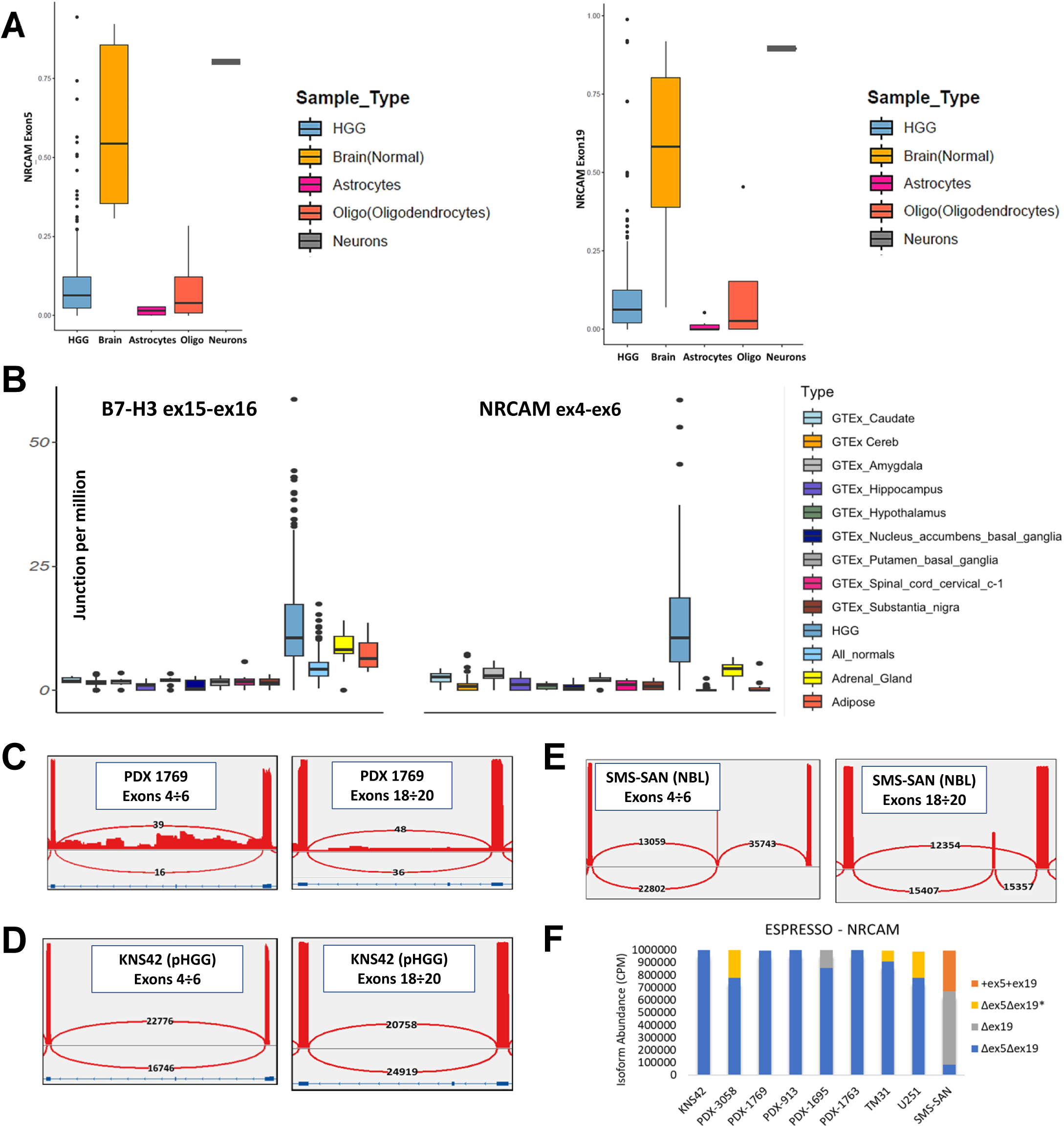
HGG-specific expression of Δex5Δex19 NRCAM isoform. **A.** Box plot showing average percent-spliced-in (PSI) value for Exon 5 and Exon 19 in NRCAM across normal brain cell types. Horizontal lines correspond to median values. **B. (Left)** Boxplots showing expression of mRNA encoding the established B7-H3 antigen in indicated GTEx tissues (neural- and non-neural) and pHGG samples. Read counts corresponding to a constitutive exon-exon junction were used to estimate transcript levels and were plotted as junctions per million counts on the *y* axis. Horizontal lines correspond to median values. **(Right)** Expression of the Δex5 isoform of NRCAM mRNA (the exon 4 – exon 6 junction) in the same samples. **C.** Validation of NRCAM exon 5 and 19 skipping in PDX7316-1769 by Direct cDNA long-read RNA-seq. Reads extending from the 3’ to the 5’ end of the NRCAM transcript were visualized in IGV and depicted as sashimi plots. **D, E.** Validation of NRCAM exon 5 and 19 skipping and inclusion in KNS42 cells and SMS-SAN cells by targeted long-read RNA-seq, respectively. **F.** Stacked plot showing estimated abundances of NRCAM Δex5 and Δex19 transcripts as measured in counts per million (*y* axis) across pHGG cell lines (KNS42) and PDXs, GBM cell lines (TM31 and U251), and a neuroblastoma cell line (SMS-SAN). ΔexΔ5ex19* represents transcripts with additional alternative splicing events besides the skipping of exons 5 and 19.

PSI values, while reflective of the underlying molecular mechanism, do not yield information about expression levels of a given isoform, which are much more relevant from the immunotherapy standpoint. Thus, we complemented our splicing analysis with the parsing of splice junction counts, one of the key outputs of the STAR aligner ^46^, as described by us recently ^47^. Specifically, we quantified expression of the non-canonical exon 4-exon 6 junctions in pHGG samples and across the GTEx dataset comprised of both neuronal and non-neuronal normal tissues. *B7-H3/CD276* canonical exon 15-exon 16 junction was used for comparison. We observed that *NRCAM* Δex5Δex19 and *B7-H3* transcripts were expressed at comparable levels in pHGG and brain sub-regions, while *NRCAM* Δex5Δex19 had the advantage of being expressed at lower levels in non-neuronal normal organs, such as the adrenal gland and the adipose tissue (Figure 2B). Similar results were obtained using as a readout *NRCAM* exon 18-exon 20 junction (Supplemental Figure S2B, top left.) The other three members of the *L1CAM* family showed the same pattern of selective exon usage, except that their expression levels in pHGG were markedly lower (Supplemental Figure S2B, top right and bottom).

To determine whether skipping of *NRCAM* microexons 5 and 19 occurs concurrently and in the context of functional, cap-to-poly(A) transcripts, we performed direct cDNA long-read sequencing using Oxford Nanopore platform PromethION P2 for two DMG PDX samples, 7316-1763 and 7316-1769. To ensure accurate detection of small exons, STAR-aligned reads were re-aligned using MisER ^48^. We observed the skipping of both *NRCAM* exon 5 and exon 19 in PDX 7316-1769 (Figure 2C), however due to low read depth we could not detect any transcripts for *NRCAM* in PDX 7316-1763. To circumvent the read depth issues, we performed targeted long-read re-sequencing on several PDX samples, KNS42 pHGG cell line, TM31 and U251 GBM cell lines, and SMS-SAN neuroblastoma cell line. Direct examination of sequencing reads in the Integrative Genomics Viewer [IGV; ^49^] revealed that all oligo(dT)-primed *NRCAM* transcripts extending to the 5’ end lacked exons 5 and 19 in KNS42 cells, but included them at ∼50% frequency in SMS-SAN NBL cells (Figure 2D,E). Using the ESPRESSO computational tool ^50^, we quantified the expression of various *NRCAM* transcripts in all sequenced samples. We observed that all pHGG and GBM cell lines and PDXs predominantly expressed the *NRCAM* variant lacking both exon 5 and exon 19 (Figure 2F, blue, grey, and yellow bars), while in SMS-SAN cells inclusion of exon 19 was common (orange bar) and skipping of both exon 5 and exon 19 was rather rare.

### Assessment of tumor heterogeneity by single-cell long-read-seq

Bulk long-read RNA sequencing is commonly used to identify full-length transcripts, but initially it did not have single-cell resolution capabilities ^51,52^. Subsequent single-cell long-read approaches were developed for fewer than 100 cells ^53,54^, making the assessment of tumor heterogeneity difficult. To overcome this limitation, we recently developed techniques for long-read sequencing of thousands of single cells from both fresh [ScISOr-Seq; ^55^] and frozen [SnISOr-Seq; ^56^] tissues. The most recent iteration of this approach [(SnISOr-Seq/ScISOr-ATAC; ^56,57^] allows for the accurate cataloging of full-length transcripts devoid of intronic sequences. We applied this approach to DMG PDX 7316-3058.

According to the short-read dataset (the 10x Genomics part of the workflow), this sample can be broken down into 3 clusters based on K-means (Figure 3A, panel i). While all of them expressed glial cell markers, proliferating cells were largely confined to Cluster 2, as evidenced by expression of the *MKI67* gene (panel ii). Nevertheless, virtually all neoplastic cells, regardless of cluster attribution, expressed both *B7-H3* (panel iii) and *NRCAM* mRNAs (panel iv), especially robustly the latter.

**Figure 3.**
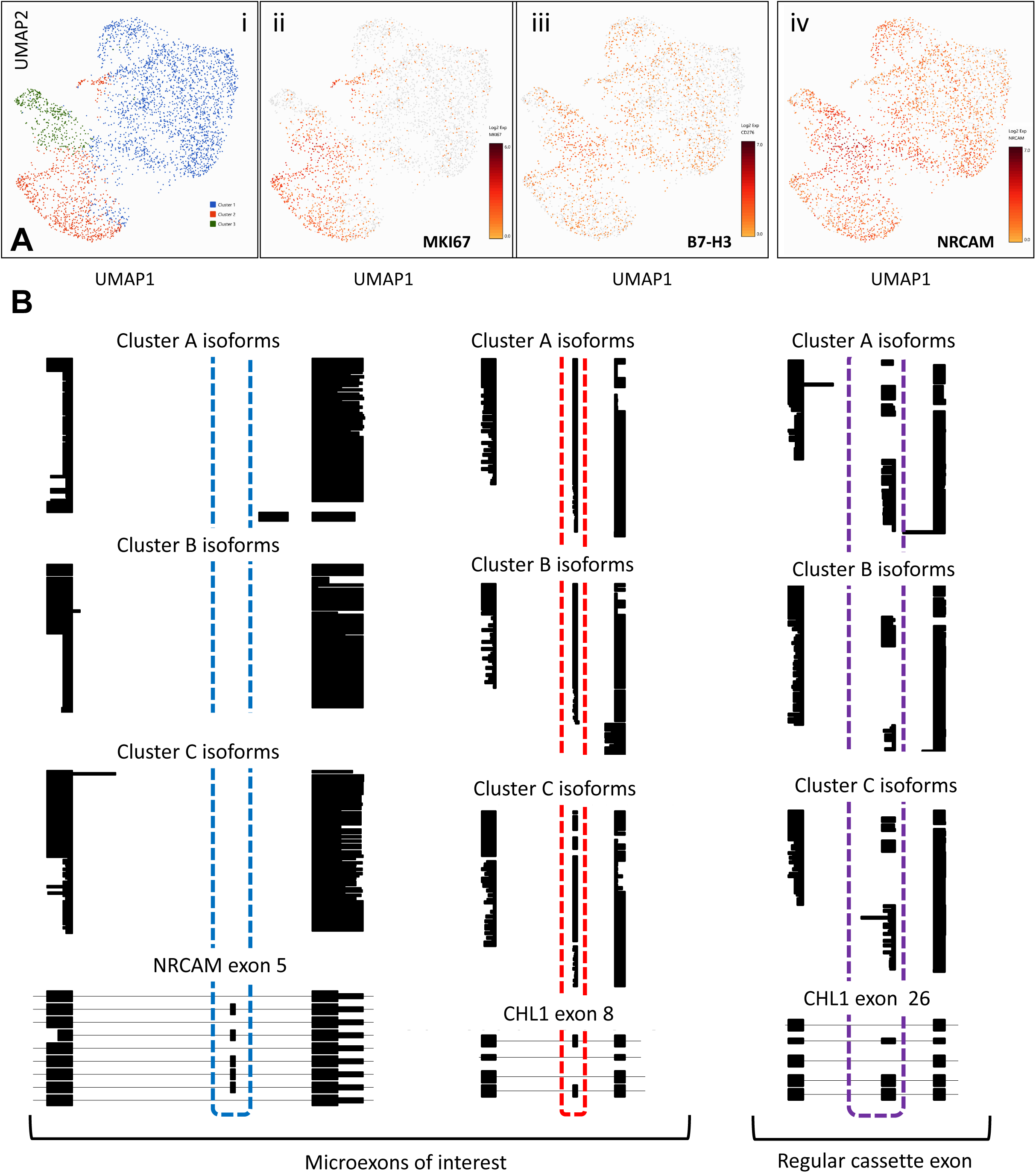
Single-nuclei analysis of PDX 7316-3058. **A.** Short-read RNA-seq analysis of >3,700 nuclei sequenced using the 10x Genomics platform. Uniform Manifold Approximation and Projection (UMAP) plots were generated using Cell Ranger and colored by cell-type annotation (i). Additional UMAP plots of the same cells were colored by expression of individual genes using Loupe Browser (ii-iv). **B.** Long-read RNA-seq analysis of full-length transcripts. ScisorWiz plots show reads spanning exons of interest in NRCAM and CHL1 genes (dotted rectangles) and grouped by clusters. Each horizontal line indicates one transcript; thick blocks denote exons, and thin lines denote introns (not drawn to scale to aid visualization). Bottom panels show annotated GENCODE transcripts.

Then, using the long-read dataset (the Oxford Nanopore part of the workflow), we asked how microexons of interest are represented in 3 major clusters (A, B, and C in Figure 3B). This was achieved by retaining barcoded reads only and aligning them over GRCh39 genome by minimap2 with gencode v44 gtf as described earlier ^58^. When we examined ScisorWiz ^56^ plots. We observed that all recovered reads for *NRCAM* exons 4 and 6 skipped microexon 5 (blue dotted rectangle) and all recovered reads for *CHL1* exons 7 and 9 included microexon 8 (red rectangle), with no evidence of intra- or inter-cluster heterogeneity. This was in sharp contrast with the *CHL1* “non-micro” cassette exon 26, whose inclusion and skipping varied considerably both within and among the 3 clusters (purple rectangle). Collectively, these results suggest that in pHGG *L1-CAM* microexons are processed in a uniform, rather than a stochastic manner, making the corresponding proteoforms much more compelling immunotherapy targets.

To validate these findings in additional pHGG samples, we parsed data from the recent study wherein bulk and single-cell RNA-Seq was performed on 19 pHGG primary samples [^59^; NCBI GEO dataset GSE231859]. Non-cancerous samples were excluded, as were samples annotated as epithelioid glioblastoma or with fewer that than 200 neoplastic cells. The bulk sequencing reads corresponding to the remaining 8 samples were visualized in IGV, with emphasis on NRCAM exons 5 and 19. As anticipated, we observed predominant patterns of exon 19 and especially exon 5 skipping, with 5 out of 8 samples not containing any exon 5 reads (Supplemental Figure S3A). Then single-cell sequencing reads were integrated and analyzed using the Seurat and Azimuth packages ^60–63^, and cell types were mapped using the Human Motor Cortex Reference Explorer. When all cell types were included in the analysis, NRCAM and B7-H3 showed similar patterns of intertumoral heterogeneity (Supplemental Figure S3B). However, after zeroing on the glial compartments (astrocytes, oligodendrocytes, and oligodendrocyte progenitor cells; Supplemental Figure S4A), we observed much more robust and more uniform distribution of NRCAM mRNAs than B7-H3 mRNAs, with fewer dropout events (Supplemental Figure S4B,C), attesting to NRCAM’s potential as an immunotherapy target.

### The role of ***Δ***ex5***Δ***ex19 NRCAM in pHGG pathogenesis

To determine the contribution of NRCAM Δex5Δex19 to gliomagenesis, we first designed a guide RNA mapping to NRCAM exon 4 (Supplemental Figure S5A) and used it in combination with the CRISPR/Cas9 system to generate *NRCAM*-null (KO) KNS42 cells. The KO event was validated at the genomic DNA levels by amplicon re-sequencing (Supplemental Figure S5B) and at the protein level - by immunoblotting with a commercially available anti-NRCAM antibody (Figure 4A). To determine whether the Δex5Δex19 NRCAM isoform is expressed on the plasma membrane, we adapted a whole-cell biotinylation assay. Briefly, live KNS42 cells were incubated with biotin to label Lys residues of surface proteins, which upon lysis were isolated using neutravidin beads and analyzed by immunoblotting. In these subcellular localization experiments, EGFR served as a reference cell surface marker and tubulin and actin as cytosolic markers. We reliably observed the Δex5Δex19 NRCAM isoform in both total cell lysate and the cell surface but not cytosolic (“flow-through”) fractions and largely absent from all NRCAM KO cells fraction (Figure 4B). This result is in line with multiple reports showing that microexon-encoded amino acids are often located on outer surfaces and regulate protein-protein interactions, as catalogued in the recent review (Table 1 in Ref. ^64^).

**Figure 4.**
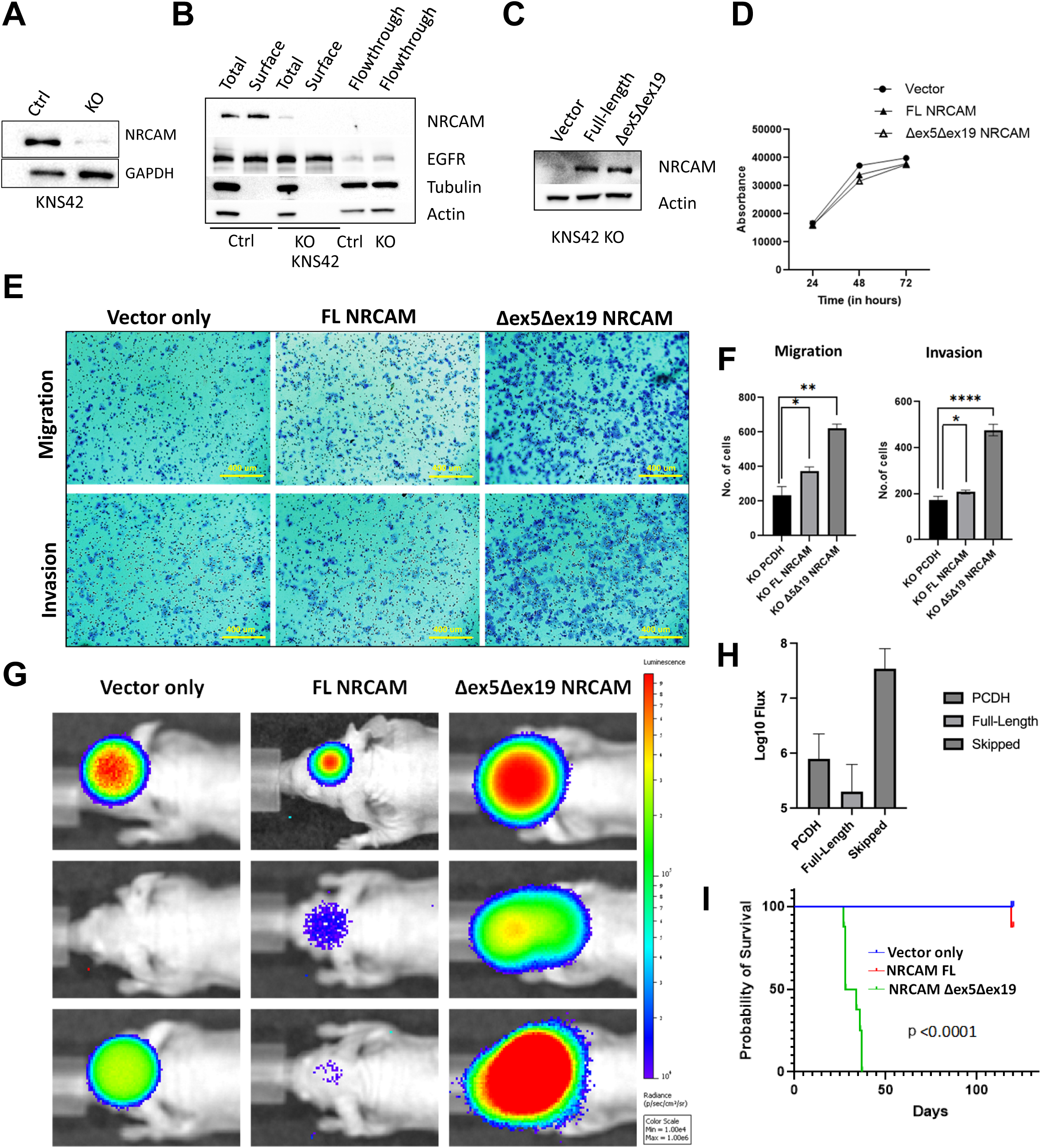
Aberrantly spliced NRCAM proteoforms in pHGG. **A.** Immunoblotting showing NRCAM levels in KNS42 pHGG cells, wild-type (Ctrl) or transfected with NRCAM-targeting gRNAs (KO). GAPDH served as loading control. **B.** Immuno-blotting showing levels of full length and Δex5Δex19 NRCAM isoforms in cell-surface and flow-through fractions, and total cell lysates of KNS42 Ctrl and NRCAM KO cells. EGFR, Tubulin and Actin were used as internal controls. **C.** Immuno-blotting showing NRCAM levels in KNS42 NRCAM KO cells reconstituted with the full length (FL) or the Δex5Δex19 NRCAM isoforms. **D.** Proliferation rates of NRCAM KO cells expressing the full length or the Δex5Δex19 NRCAM isoforms. **E.** Migration (top) and invasion (bottom) potential of NRCAM KO cells re-expressing full length and Δex5Δex19 isoforms of NRCAM. The Transwell assays were performed without or with Matrigel layers, respectively. **F.** Quantitation of migrated and invaded cells from G. Each bar represents mean ± SEM. Significance (asterisks) was determined using unpaired Student’s t-test. **G.** Optical imaging of the same cells additionally engineered to expressed firefly luciferase and orthotopically injected into the cortices of NSG mice. 3 mice out of 8 were imaged on day 28. **H.** Raw photon counts corresponding to tumors in panel G. Each bar represents mean ± SEM. **I.** Kaplan-Meier curves representing survival of mice depicted in panel G, with p value determined by log-rank (Mantel-Cox) test.

To determine the role of the skipped isoform of NRCAM in pHGG, we reconstituted NRCAM KO cells with the full-length (exon 5-/exon 19-including) and the Δex5Δex19 NRCAM isoform and confirmed their expression by immunoblotting (Figure 4C). Compared to vector control cells, neither of them showed any effect on cell proliferation in 2-D cultures (Figure 4D). However, the Δex5Δex19 - but not the full-length NRCAM isoform - enhanced migration and invasion by KNS42 cells, as evidenced by the Transwell assays performed without and with Matrigel layers, respectively (Figure 4E, F), as described by us previously ^65^.

Finally, we modified these cells to express firefly luciferase and implanted them orthotopically into the cortexes of non-obese-diabetic severe combined immunodeficiency gamma (NSG) mice, as described previously ^66^. We observed more consistent tumor take and substantially more robust tumor growth when the Δex5Δex19 NRCAM isoform was expressed, whereas both *NRCAM*-null and *NRCAM*-full length KNS42 cells established and grew rather poorly (Figure 4G,H). In fact, mice bearing these tumors were still alive on Day 120, while their Δex5Δex19 counterparts were all dead by day 30 (Figure 4I). The apparent essentiality of Δex5Δex19 for tumor growth and its uniform expression on the cell surface made it an even more attractive target for pHGG immunotherapy.

### pHGG-selective anti-NRCAM antibodies and their therapeutic utility

Given the minimal difference in the amino acid sequences between the full-length and the Δex5Δex19 NRCAM proteoforms, we used AlphaFold 3 ^67^ to predict 3D structural models for the corresponding UniProt-annotated ectodomains. It predicted a “horseshoe” conformation with a hinge between Ig-like 2 and Ig-like 3 domains in both isoforms, similar to crystal structures of another L1 family-member, L1CAM [^68^; discussed in ^69^]. AlphaFold 3 also predicted that the proline-rich microexon 19-encoded amino acid sequence creates a sharp bend between Ig-like 6 domain and the first fibronectin type-III domain (Figure 5A, left). This bend is absent from the NRCAMΔex5Δex19 proteoform model, creating a more open conformation that could be available for selective targeting (Figure 5A, right).

**Figure 5.**
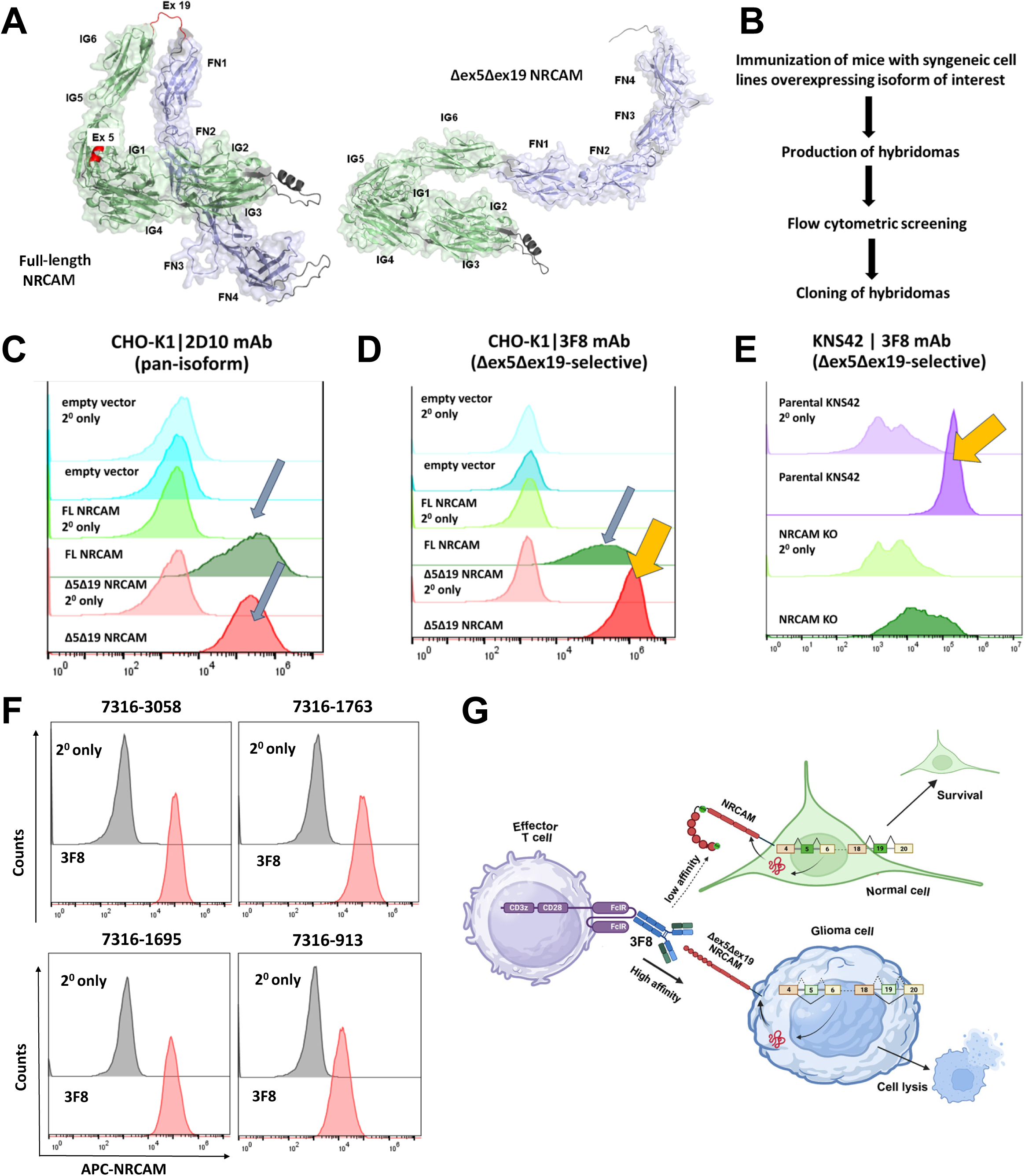
Detection of NRCAM proteoforms by mAb 3F8. **A.** AlphaFold models of the ectodomains of the canonical (left) and Δex5Δex19 (right) NRCAM isoforms, with the signal peptide (amino acids 1-24) removed. Ectodomain and signal peptide were identified using UniProt annotations. Ig-like domains are shown in green, fibronectin type III domains in purple, and microexons 5 and 19 in red. **B.** Schematic showing the pipeline followed for antibody production**. C.** Flow cytometry histograms showing 2D10 and **D.** 3F8 mAb binding profiles when used on live CHO-K1 cells expressing “empty vector” (target-null, blue), full-length (FL) NRCAM (unintended target, green), and Δex5Δex19 NRCAM (intended target, red). Control staining’s with secondary (2^0^) antibody only are shown for comparison. **E.** Same staining performed on live HGG KNS42 cells endogenously expressing Δex5Δex19 NRCAM (intended target, purple) or with the entire gene knocked out using CRISPR-Cas9 (target-null, green). In both panels, orange arrow points to the signal generated by Δex5Δex19 NRCAM. **F.** Flow cytometry histograms showing 3F8 binding to various patient-derived pHGG cells compared to secondary-antibody-only stained samples (2°). **G.** The strategy to test the therapeutic utility of the 3F8 antibody against glioma cells expressing the splice isoform of NRCAM. The composition of FcγRI-based UIR is shown on the left. Non-neoplastic cells expressing the full-length isoform of NRCAM are depicted at the top right as being presumably resistant to 3F8 UIR treatment.

To generate NRCAM Δex5Δex19-selective monoclonal antibodies, we expressed this isoform in murine NIH3T3 cells and immunized syngeneic C57BL/J mice with whole cells using the Fred Hutchinson Cancer Center Antibody Technology Core Facility (Figure 5B), as described by others previously ^70^. Supernatants from the resultant hybridomas were used to stain CHO cells expressing either the full-length or the Δex5Δex19 NRCAM proteoform. As expected, the majority of mAbs (exemplified by 2D10 in Figure 5C) stained both the full-length and the NRCAM isoforms equally well (grey arrows). However, mAb 3F8 showed selectivity for NRCAM Δex5Δex19, as judged by ∼10-fold difference in mean fluorescent intensities (Figure 5D, grey and orange arrows). Importantly, this binder recognized the endogenously expressed Δex5Δex19 NRCAM in pHGG samples, such as the KNS42 cell line (Figure 5E, purple peak with orange arrow), even though non-specific, background staining was also observed in *NRCAM* KO cells (green peak). We also stained 4 pHGG PDXs with 3F8. Consistent with single-cell RNA-seq data (Supplemental Figures S3 and S4), we observed robust and uniform expression of NRCAM Δex5Δex19 (Figure 5F).

To determine how efficient and selective 3F8-based immunotherapeutics might be, we applied a universal immune receptor (UIR) technology to redirect T cell specificity towards antibody-stained cells, as described by us previously ^71–73^. Specifically, we “painted” KNS42 cells with the 3F8 antibody and admixed them in vitro with human donor-derived T cells engineered to express a CD64 (Fc gamma receptor I; FcγRI)-based UIR (Figure 5G). We observed that this antibody-UIR combination could kill the majority of pHGG cells, but only when the cancer cells expressed the Δex5Δex19 NRCAM proteoform, ectopically or endogenously (KNS42 or PDX 7316-3058) (Figure 6A, top row). No killing was observed against cells expressing full-length NRCAM, or when the FcγRI UIR was armed by an irrelevant isotype-control IgG2b antibody (Figure 6A, bottom row), or using un-transduced donor T cells (Supplemental Figure S6A). We further tested the 3F8-UIR combination against adult GBM cell lines U251 and TM31 (which exclusively express the Δex5Δex19 NRCAM isoform) and their *NRCAM* KO derivatives. We observed the same efficacy and selectivity of cell killing as seen with pHGG models (Figure 6B and Supplemental Figure S6B). We thus concluded that mAb 3F8 is a promising immunotherapeutic for pediatric and adult tumors of glial origin.

**Figure 6.**
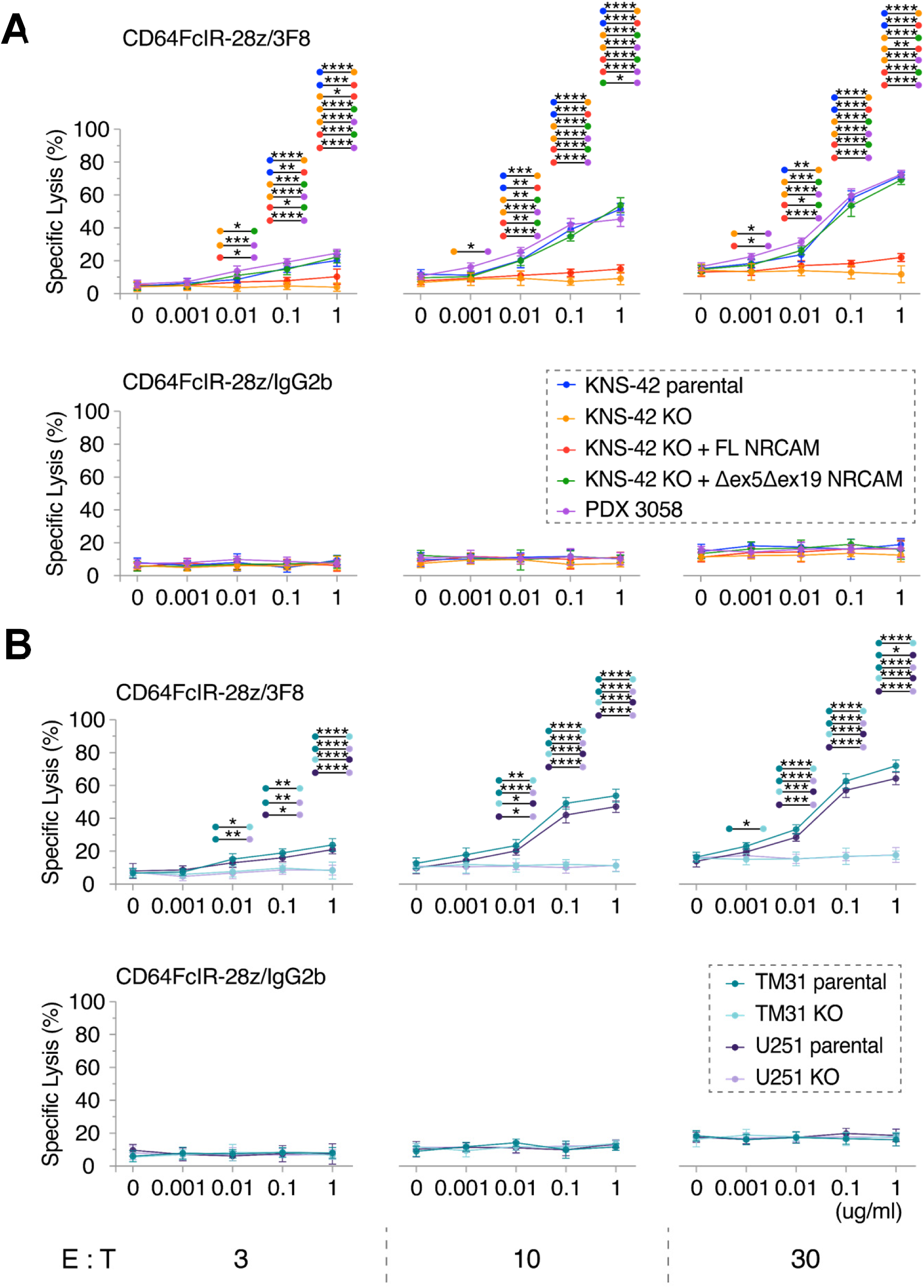
mAb 3F8-mediated killing of glioma cells. **A.** Killing of PDX3058 and KNS42 cells expressing indicated NRCAM isoforms. **B.** Killing of adult glioblastoma U251 and TM31 cells and their NRCAM KO derivatives. In both panels, shown on the *x* axis are Ab concentrations (in mg/ml), and on the *y* axis - the extent of tumor cell killing, as evidenced by reduced luciferase expression. “E:T” values refer to the ratio of effector (T) to target (glioma) cells. Two-way Anova was used for comparison across different treatment conditions and groups. **** - p<0.00001, *** - p<0.001, ** - p<0.01, * - p<0.05

## DISCUSSION

It is well-recognized that failures of adoptive cancer immunotherapies come in two flavors: effector cell-centric (e.g., T cell exhaustion) and target cell-centric (e.g., epitope loss or down-modulation) ^74^. The latter problem underscores the need for robust pipelines yielding alternative antigens for salvage immunotherapies. Traditionally, such pipelines focused either on discovery of lineage-specific markers (B cell-specific CD19, CD22, etc.) ^75^ or on identification of proteins overexpressed in tumors compared to their respective tissues of origin. While such “tumor-versus-normal” comparisons continue to yield viable therapeutic targets, for example DLK1 and GPC2 in high-risk neuroblastoma ^76,77^, GPC2 in CNS tumors ^78^, B7-H3 (CD276) and GD2 in diffuse intrinsic pontine glioma (DIPG) and DMG ^15,20^, GD2 in medulloblastoma ^79^, this approach might be entering the phase of diminishing returns, as existing microarray and RNA-seq datasets have been mined extensively already. Additionally, “tumor-versus-normal” comparisons often ignore expression of the target antigen in non-adjacent normal tissues, creating the potential for severe on-target, off-tumor toxicities (OTOT) ^80^. For CAR T cells, OTOT could be circumvented with locoregional delivery, but this approach may not be effective for antibody-drug conjugates and other soluble therapeutics.

All these limitations led the cancer immunotherapy field to look for alternative sources of tumor-specific epitopes, such as alternatively splicing ^81,82^. In a hallmark paper investigating splicing patterns across The Cancer Genome Atlas, it was estimated that in the average tumor there exist close to 1,000 unique “neojunctions” not typically found in normal samples profiled by the GTEx consortium ^83^. However, only a tiny fraction of them (∼0.1%) generate peptides capable of being presented by major histocompatibility complex class I molecules.

In principle, this bottleneck could be bypassed by focusing on transmembrane proteins with large extracellular domains, which could be recognized by engineered T-cells independently of MHC class I presentation ^84^. However, it is not clear what fraction of alternatively spliced transcripts yields functional proteoforms able to reach the plasma membrane. In hematologic malignancies, for example, there are several examples where exon skipping results in either protein degradation [P2RX5 ^85^] or its retention in the endoplasmic reticulum [CD19 ^86,87^, CD33 ^88^]. Finally, even when alternative splicing-derived proteoforms are expressed on the cell surface, anti-peptide monoclonal antibodies directed against “neojunctions” might efficiently and specifically recognize the denatured protein [e.g., CD22 Δex5-6 ^89^], but not necessarily its membrane-bound conformation. Thus, in most cases alternative splicing might be just as likely to drive resistance than to enable new therapies ^90^. However, not all alternative splicing events are created equal; and some of them, for example inclusion/skipping of microexons, have evolved to preserve protein structures while generating unique 3D conformations ^29,64^.

Here we report that in pediatric HGG, the pattern of microexon inclusion is markedly different from that observed in normal brain samples. In fact, microexon skipping was the only SE type of event with profound differences in ΔPSI values (>50%). By focusing further on SE events with high prevalence, we were able to identify members of the L1 family of cell adhesion molecules ^69^, including NRCAM, as being particularly strongly affected by alternative splicing. This is not completely surprising since NRCAM was recently shown to be the molecule with the greatest number of proteoforms in the developing mouse and human retinas ^91^. There is also some evidence that NRCAM and related CAMs might be implicated in tumor progression [reviewed in ^92^]. However, the importance of individual proteoforms in either normal or development and cancerous growth has not been elucidated.

By focusing on concurrent skipping of *NRCAM* microexons 5 and 19, we were able to demonstrate that not only the corresponding proteoform is stable and present on pHGG cell surfaces, but it is also essential for cell migration and invasion in vitro and for progressive tumor growth in orthotopic mouse models. Remarkably, despite minimal amino acid changes, it is also antigenically distinct, as we were successful in generating a monoclonal antibody (3F8) with high selectively toward NRCAM Δex5Δex19. This selectivity manifested itself not only in flow cytometry experiments, but also in functional cell killing assays, where pHGG cells reconstituted with NRCAM Δex5Δex19 – but not the full-length isoform – were effectively killed by human T cells engineered to express an FcγRI-based universal immune receptor and redirected by an NRCAM Δex5Δex19-specific mAb. These findings provide the rationale for future development and preclinical testing of other 3F8-based immunotherapeutics, including CARs.

If 3F8-based immunotherapeutics prove to be as potent as current ones (e.g., B7-H3 CARs), targeting NRCAM Δex5Δex19 could be advantageous from the OTOT standpoint. Indeed, we observed that this mRNA isoform is expressed at lower levels in several normal tissues, including the adrenal gland. Even in normal brain samples, which have sizable fractions of cells of glial origin, we failed to detect significant levels of *NRCAM* Δex5Δex19 expression, as judged by JPM counts (Figure 2B). In fact, the recently developed Cancer-Specific Exon Miner identified NRCAM as one of the six alternatively spliced proteoforms highly and specifically expressed in multiple solid and brain pediatric tumors with high prevalence ^93^. Our data indicate that the potential of NRCAM Δex5Δex19 as a therapeutic target is not limited to pediatric tumors and could include adult low-grade gliomas and GBMs, pheochromocytomas and paragangliomas, and also cancers of non-neural origins, such lung adenocarcinomas.

The rather broad spectrum of *NRCAM* Δex5Δex19 expression raises interesting questions about its contribution to neoplastic growth. It could be argued that the skipping of microexons 5 and 19 in pHGGs simply reflects their glial, not neuronal origin; and that normal glia (and by extension, its malignant counterparts) simply don’t express the RNA-binding protein machinery necessary to include microexons. While there is some recent experimental and computational support for this “skipping by neglect” scenario ^94^, several lines of evidence indicate that it may not be universally applicable. First and foremost, the identity of cell(s) of origin for human HGG is often inferred from mouse lineage tracing experiments and therefore is not firmly established ^95^. Examination of molecular signatures frequently points to the role of oligodendrocytic progenitor cells (OPC), at least in certain anatomic locations, such as pons. In contrast, diffuse hemispheric gliomas (of which KNS42 is representative) are now thought to arise from interneuronal precursors ^96^. Second, even if most pHGG do arise from OPCs, glia-specific microexons also have been identified ^97^. Third, intermediate levels of *NRCAM* exon 19 (but not exon 5) inclusion in pheochromocytomas and paragangliomas revealed by MAJIQlopedia argue that these cassettes are dynamically and independently regulated to shape the neoplastic phenotypes of individual tumor types.

For example, skipping of *NRCAM* microexons could regulate its interactions with other members of the L1-IgCAMs ^69,98,99^ and the metastasis-promoting proteins of the ERM family [ezrin/radixin/moesin; ^100^]. Elucidation of these interactions could reveal additional opportunities for the targeting of alternative splicing in pediatric gliomas.

### Limitations of the study

Our conclusion that NRCAM Δex5Δex19 is an attractive target for adoptive T cell therapy is based largely on our success in generating a splice isoform-selective antibody (3F8) and its potency in the in vitro killing assays using UIR-armed T donor cells. We are yet to test these UIR T cells in vivo. Nor have we evaluated cross-reactivity of 3F8 with normal tissues, given the challenges of performing immunohistochemistry with antibodies recognizing conformational epitopes.

## RESOURCE AVAILABILITY

### Lead contact

Further information and requests for reagents may be directed to and will be fulfilled by the lead contact Andrei Thomas-Tikhonenko.

### Materials availability

- pHGG specimens used in this study are available upon request from the Children’s Brain Tumor Network
- The 3F8 hybridoma will be available upon request following publication of the patent application.

### Data and code availability

- Newly generated single-cell RNA-seq data have been deposited in GEO: https://www.ncbi.nlm.nih.gov/geo/query/acc.cgi?acc=GSE301658. Access to existing genomic files from CBTN is enabled through KidsFirst Portal: https://portal.kidsfirstdrc.org. Access to already processed genomic data is enabled through PedcBioPortal: https://pedcbioportal.kidsfirstdrc.org. Controlled access CBTN data were retrieved through dbGaP Study phs002517.v3.p2. The data from this study were made available pre-publication without embargo to support rapid and collaborative research in pediatric cancer via the NCI’s Cancer Research Data Commons (https://datacommons.cancer.gov). This availability was made possible with the support of NCI’s Childhood Cancer Data Initiative and Gabriella Miller Kids First Pediatric Research Program. Controlled access GTEx data were retrieved through dbGaP Study phs000424.v10.p2. The Genotype-Tissue Expression (GTEx) Project was supported by the Common Fund of the Office of the Director of the National Institutes of Health (https://commonfund.nih.gov/GTEx). Additional funds were provided by the NCI, NHGRI, NHLBI, NIDA, NIMH, and NINDS.
- This paper does not report any original code. ScisorWiz, a Linux-based R-package for visualizing differential isoform expression, had been deposited on GitHub previously: https://github.com/ans4013/ScisorWiz.
- Original western blot images have been deposited at Mendeley (https://10.17632.cmsfkvw76m.2). Microphotographs and any additional information required to reanalyze the data reported in this paper are available upon request from the lead contact.

## ACKNOWLEDGMENTS

This research has been supported by NIH grants U01 CA232563 and R03 CA293992 (to ATT), R03 DE033366 (to JLR), R01 HG013359 (to KW), and UG3 CA290451 and RO1 EB026892 (to DJP Jr). Key support was provided by the Cure Search for Children’s Cancer Foundation Acceleration Initiative Award (to ATT), and also by Cancer Research Society Next Generation of Scientists Award (to MQV), the Children’s Brain Tumor Network, and the Chad Tough Foundation. JL further acknowledges support by the National Science Foundation Graduate Research Fellowship Program (NSF-GRFP, grant DGE-2236662). ATT is Mildred L. Roeckle Endowed Chair in Pathology at Children’s Hospital of Philadelphia and Investigator on the St. Baldrick’s Foundation EPICC Team. Some schematics in the manuscript and the graphical abstract have been created with Biorender.com

## AUTHOR CONTRIBUTIONS

Conceptualization, P.S., A.S.N., J.L.R., A.C.R. and A.T.T.; methodology, A.C., Z.A., M.Q.V., K.W., A.F., Y.B., J.B.S., H.U.T., T.D., and D.J.P.; experimentation, P.S., M.H., J.L., K.H., J.J., J.F., J.D., and A.M.; data acquisition and analysis, A.S.N., T.K., B.M., P.M., G.W., K.E.H., C.D., A.F., M.Q.V., writing—original draft, P.S. and A.T.T.; writing—review & editing, P.S., G.W., and A.T.T.; funding acquisition, J.B.S., Y.B., and A.T.T.; resources, D.M., A.F., A.C.R.; supervision, J.L.R, H.U.T., T.D., D.J.P., and A.T.T.

## DECLARATION OF GENERATIVE AI AND AI-ASSISTED TECHNOLOGIES

During the preparation of this work, no generative AI or AI-assisted technologies have been used.

## STAR□METHODS

### KEY RESOURCES TABLE

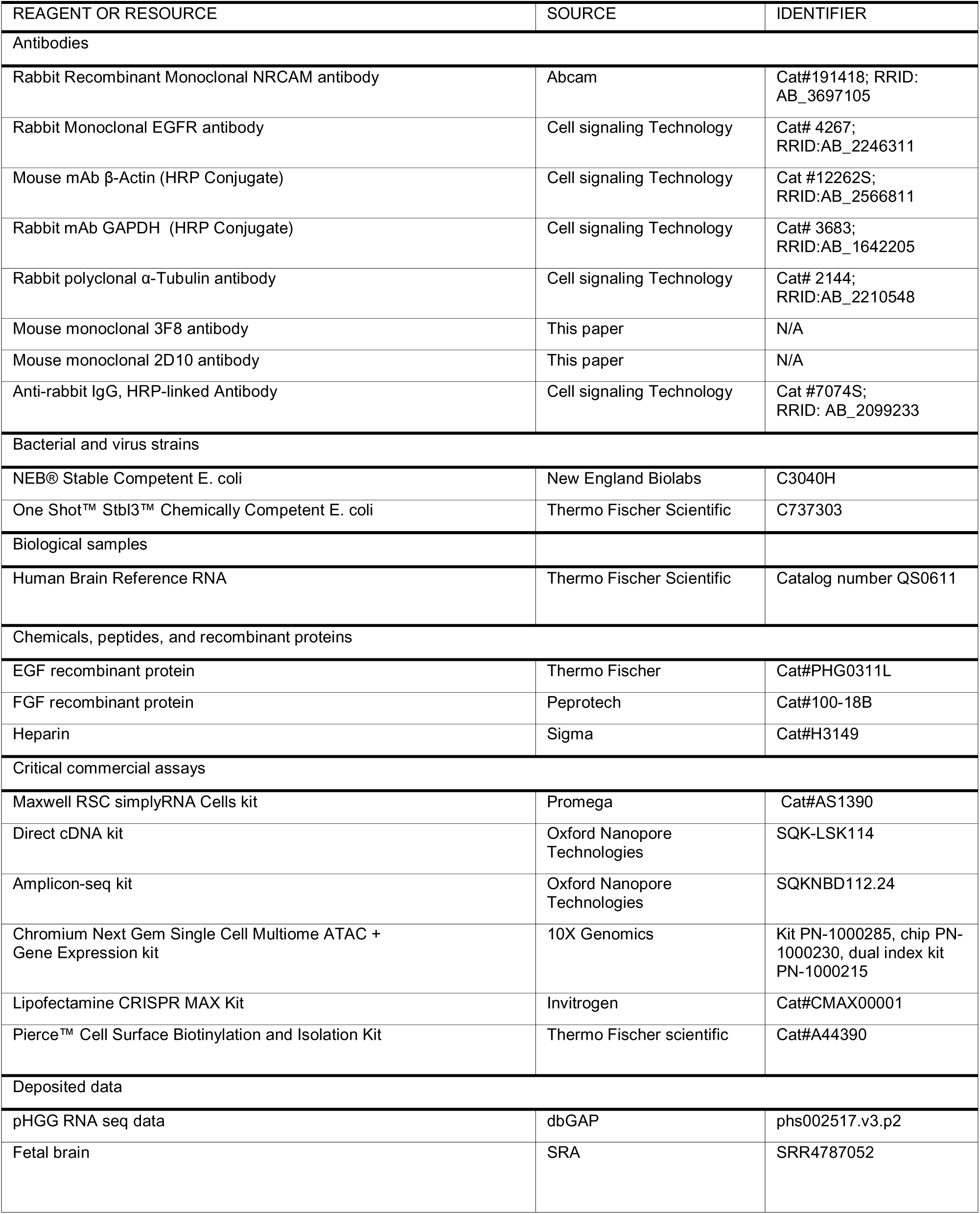
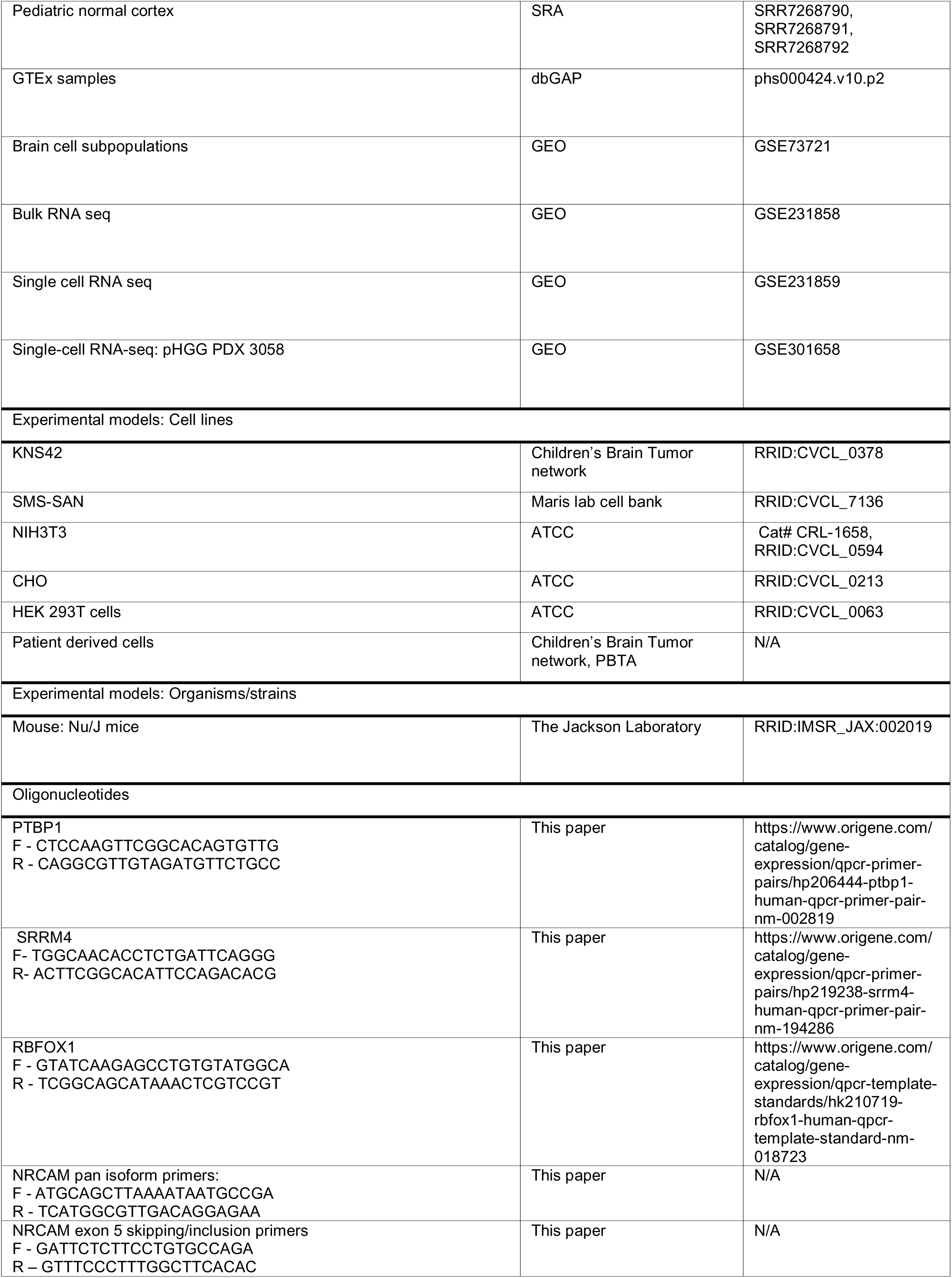
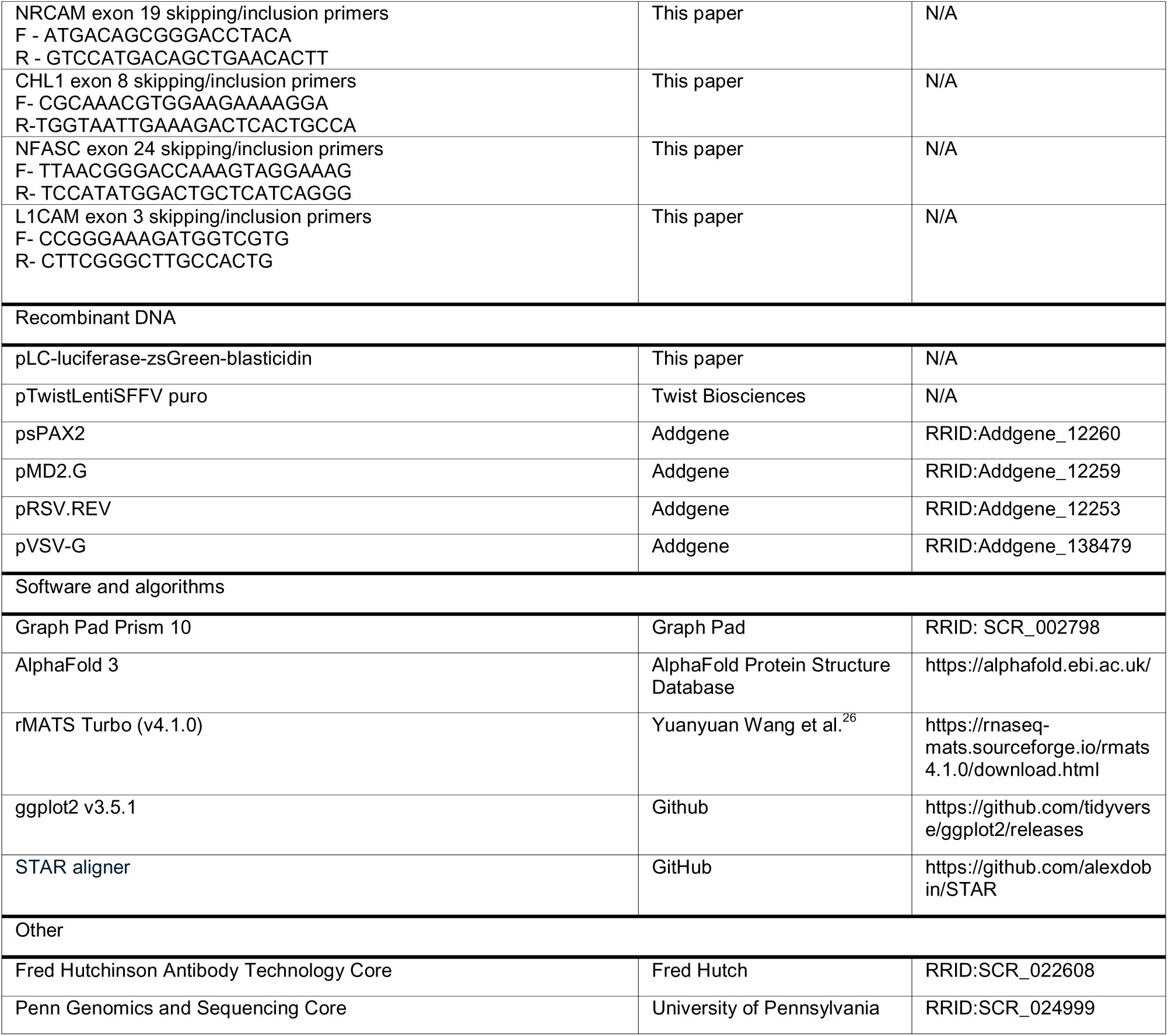

## EXPERIMENTAL MODEL AND STUDY PARTICIPANT DETAILS

### Animal experiments

All animal experiments had received prior approval from the Children Hospital of Philadelphia Institutional Animal Care and Use Committee (IACUC) and were conducted as described in Protocol IAC 24-001295 “Validation of drivers in tumor formation and preclinical testing” (approval date 2/29/2024). The orthotopic pHGG models have been established as follows. Eight to 10-week-old Nu/J mice (8 animals per treatment group) were anesthetized with 2.5% isoflurane, placed into the stereotactic frame, then injected with 1x106 glioma cells. Stereotactic coordinates used to target the cortex for tumor were as follows: -0.4 mm posterior to bregma, 3mm lateral to bregma, and 1 mm depth from pia. All animals were subjected to regular optical imaging beginning at 3 weeks post-injection.

### Human samples

All high-grade glioma pediatric brain tumor raw data were harvested from the database of Genotypes and Phenotypes (dbGAP) (accession number phs002517.v2.p2), which includes tumors from the Children’s Brain Tumor Network (https://cbtn.org) and the Pediatric Neuro-Oncology Consortium (https://pnoc.us/).

### Cell culture

The pediatric high-grade glioma cell line KNS42 was cultured in DMEM-F12 (GIBCO Cat#11320033), the neuroblastoma cell line SMS-SAN was cultured in RPMI (Corning Cat#10-040-CM), murine NIH3T3 cells were cultured in IMDM (ThermoFischer Scientific Cat#12440053), Chinese Hamster Ovary (CHO) cells were cultured in Ham’s F 12K (Kaighn’s) Medium (ThermoFischer Scientific Cat# 21127022) supplemented with 10% FBS (GIBCO Cat#26140079), 2 mmol/L L-glutamine (GIBCO Cat#25030081), and penicillin/streptomycin (GIBCO Cat #15140122) at 370C and 5% CO2. After thawing, cells were authenticated by short tandem repeat analysis, tested for Mycoplasma using the EZ-PCR Mycoplasma Detection Kit (Biological Industries Cat#20-700-20), and used for up to 12 passages. Patient derived cells were cultured in DMEM-F12 supplemented with B27 supplement (ThermoFischer Cat#12587-010), N2 supplement (ThermoFischer Cat#175020-48), EGF recombinant protein (ThermoFischer Cat#PHG0311L), FGF recombinant protein (Peprotech Cat#100-18B), Heparin (Sigma Cat#H3149), 2 mmol/L L-glutamine, and penicillin/streptomycin at 370C and 5% CO2. supplemented with 10% FBS, 2 mmol/L L-glutamine and penicillin/streptomycin at 370C and 5% CO2.

## METHOD DETAILS

### RNA extraction and reverse transcription

Total RNAs were isolated using Maxwell RSC simplyRNA Cells kit with the Maxwell RSC48 Instrument (Promega Cat#AS1390) and reverse-transcribed using SuperScript IV (Invitrogen Cat#18090010). Primer sequences used for cDNA amplification are listed in the Key resource table.

### Oxford Nanopore Technologies (ONT) long-read direct cDNA sequencing

Total RNAs were isolated using Maxwell RSC simplyRNA Cell Kits (Promega Cat#AS1390). Approximately 500 ng of total RNA was used for direct cDNA (SQK-LSK114; Oxford Nanopore Technologies [ONT]) library preparation. Subsequently, each library was loaded into a PromethION Flow Cell R10.4.1 version (FLO-PRO114M; ONT) and sequenced using a PromethION 2 Solo (ONT) for 48 hours. Raw Fast5 files were converted to fastq with guppy (version 3.4.5), followed by alignment to the GENCODE version of hg38 (version 30) using minimap2 (version 2.18); the resulting bam file was visualized using the Integrative Genomics Viewer (version 2.11.0).

### ONT targeted resequencing

For NRCAM, primers were designed to bind to exons present in all the isoforms to ensure full coverage of all alternative splicing events. 5ng of cDNA were amplified with the long LongAmp Taq 2X Master Mix (New England Biolabs) for 25 cycles. The resulting amplicons were subjected to amplicon-seq (SQKNBD112.24, ONT) library preparation. Subsequently, each library was loaded into a Spot-ON flow cell R9 Version (FLO-MIN112, ONT) and sequenced in a MinION Mk1C device (ONT) until it had acquired at least 1000 reads per sample. Acquired reads were aligned using Minimap2 version 2.24-r1122 and visualized in IGV version 2.12.3. When indicated, MisER, a splicing re-alignment tool, was run on ONT output files and reads were realigned using NRCAM RefSeq transcripts downloaded from UCSC genome browser. The flanking region parameter for Miser was 110. For isoform quantitation, ESPRESSO was run with default settings on MisER-realigned bam files.

### Single-nuclei isoform RNA sequencing

Nuclei from pHGG patient-derived organoids were isolated and processed using the Chromium Next Gem Single Cell Multiome ATAC + Gene Expression kit from 10X Genomics (kit PN-1000285, chip PN-1000230, dual index kit PN-1000215). For short-read sequencing, the Chromium Single Cell Multiome Gene Expression Library was performed by following the manufacturer’s instructions, starting from 8,000 single-nuclei. It was then loaded on an Illumina NovaSeq 6000 with PE 2 × 100 paired-end kit, using the following sequencing read lengths: 28 cycles Read1, 10 cycles i7 index, 10 cycles i7 index, and 90 cycles Read2. For long-read sequencing, cDNA obtained from the 10X Genomics Single Cell Multiome ATAC + Gene Expression kit was used for amplification and exome enrichment. The amplified/enriched cDNA was then sequenced on the Oxford Nanopore Technology platform starting from ∼200 fmol cDNA and using the Ligation Sequencing Kit (SQK-LSK114), according to the manufacturer’s protocol (Nanopore Protocol, Amplicons by Ligation, version ACDE_9163_v114_revO_29Jun2022). The ONT library was loaded onto a PromethION sequencer by using PromethION Flow Cell (FLO-PRO114M) and sequenced for 72□h.

Base-calling was performed with Guppy by setting the base quality score >7.

### Genome editing

Single-guide RNA targeting NRCAM exon 4 and CAS9 protein were obtained from Integrated DNA Technologies. Cas9 ribonucleoprotein complexes were assembled following manufacturer’s recommendations. These ribonucleoprotein complexes were transfected into KNS42 and 7316-3058 cells using Lipofectamine CRISPR MAX Kit (Invitrogen Cat#CMAX00001) following manufacturer’s instructions. The effect of gRNA was determined by immunoblotting for NRCAM.

### Plasmid constructs and viral infections

For NRCAM constructs, the coding sequence of the full-length isoform (NM_001193582.2/ENST00000413765.6) and Δex5Δex19 NRCAM (NM_005010.5/ENST00000351718.8) was synthesized and cloned into the lentiviral backbone (pTwistLentiSFFV puro) by Twist Biosciences. For viral particle production, 293T cells (ATCC) grown to 90% confluence in a 10-cm Petri dish were transfected with 10 μg of the NRCAM constructs, 2.5 μg of the pMD2.G (RRID:Addgene_12259) and 7.5 μg of the psPAX2, (RRID:Addgene_12260) using Lipofectamine 3000 (Invitrogen, #L3000001). Supernatants were collected 48 hours after transfection, passed through 0.45-μm PVDF filters, and incubated for 48 hours with cells in media containing 10 µg/mL polybrene (MedChemExpress, Cat#HY112735). After 48 hours, culture media were replaced, and cells were incubated with 1.5 μg/mL puromycin for selection. For FcIR-28z, transfection was performed with a plasmid mix containing the transfer plasmid encoding the Fc binding immune receptor (FcIR) fused to the intracellular TCR and co-stimulatory signaling domains (FcIR-28z), along with packaging plasmids pRSV.REV (RRID:Addgene_12253), pMD2.G, and pVSV-G (RRID:Addgene_138479) in a 15:18:18:7 mass ratio. Lentiviral supernatants were collected at 24- and 48-hours post-transfection, filtered through a 0.45 µm filter, and concentrated via ultracentrifugation at 25,000 rpm for 2.5 hours. The final viral preparation was stored at -80°C. Viral titers were determined by transducing HEK 293T cells, and results were expressed as Infection Units per milliliter (IU/mL).

### Immunoblotting

Total cell lysates were prepared from cells using RIPA buffer with protease and phosphatase inhibitors (Pierce Halt Inhibitor Cocktail, Thermo Fisher Scientific Cat#78446). After protein transfer to PVDF (Immobilin-p, Millipore Cat#IPVH00010), anti-NRCAM antibody (Abcam Cat#191418, RRID: AB_3697105) and anti-EGFR antibody (Cell Signaling Technology Cat# 4267, RRID:AB_2246311) were used. Subsequently, recommended dilutions of horseradish peroxidase–conjugated secondary antibody (Cell signaling, #7074S, RRID:AB_2099233) were added. Enhanced chemiluminescence (Millipore, #WBKLS0500) was used to detect bands that were then captured by Chemiluminescence imager (GE Healthcare). For internal normalization, HRP-conjugated antibodies against β-actin (Cell Signaling Technology Cat #12262S, RRID:AB_2566811), GAPDH (Cell Signaling Technology Cat# 3683, RRID:AB_1642205), or tubulin (Cell Signaling Technology Cat# 2144, RRID:AB_2210548) were used.

### Isolation of Biotin-labeled Cell Surface Proteins

Cell surface biotinylation was performed using Pierce™ Cell Surface Biotinylation and Isolation Kit (Thermo Fischer scientific Cat#A44390). Briefly, cells were grown to confluency in 10-well dishes, washed with PBS, and labeled with a non-cell-permeable sulfo-NHS biotin analog for 10 minutes at room temperature. After washing, ice cold TBS was added to the cells which were collected with a cell scraper, and pelleted by centrifugation (500 × g, 5 min, 4 °C). To isolate biotin-labeled cell surface proteins, lysates were loaded into Neutravidin agarose columns and incubated for 30 minutes at room temperature. Eluted protein fractions were then analyzed by immunoblotting.

### Cell proliferation, migration, and invasion assays

To assess proliferation, cells were seeded in white 96 well flat bottom plates at a density of 8000 cells per well. Cell viability was measured using the CellTiter-Glo (CTG) luminescent assay (Promega Cat#G7570). Luminescence was measured using Biotek Synergy 2 plate reader at 24, 48 and 72hrs. Corning BioCoat Cell Culture Inserts and Matrigel inserts (Corning, USA) were used for migration and invasion assays, respectively, as described in (58). Briefly, the inserts were rehydrated with plain DME-F12 for 2 h before use. 5–7×104 cells pre-treated with 10ug/ml of Mitomycin C were trypsinized and resuspended in serum-free medium and then seeded onto 24-well Transwell chambers with 8-μm pore membrane in 500 μL serum-free medium. The lower chamber contained medium supplemented with 10% FBS. After incubation for 22 h, the non-migrated/invaded cells on the upper side of membrane were removed with a cotton swab and the migrated/invaded cells stained with crystal violet and microphotographed.

### AlphaFold analysis of NRCAM isoform structures

AlphaFold 3 models were generated for the UniProt-annotated ectodomains of the full-length and Δex5Δex19 NRCAM isoforms, specifically C9JYY6 and Q14CA1. The signal peptide at the N-terminus (amino acids 1-24, per UniProt) was removed before generating the models. Additionally, these isoforms lack the fifth fibronectin type-III domain, owning the exonic structures of the respective mRNAs (ENST00000413765.6 and ENST00000351718.8).

### Generation of monoclonal antibody against NRCAM ***Δ***ex5***Δ***ex19

All monoclonal antibodies were generated at the Fred Hutchinson Antibody Technology Core. Briefly, male and female 20-week-old mice were immunized with syngeneic NIH3T3 cells overexpressing Δex5Δex19 NRCAM. Following a 12+ week boosting protocol, splenocytes were isolated from high-titer-yielding mice and electrofused with FOX-NY myeloma cells. Hybridomas secreting isoform specific antibody were identified and isolated. Antibodies from the picked clones were validated for proteoform binding by flow cytometry using cell-based flow cytometry. Clone 3F8 was subcloned followed by validation for proteoform binding by cell-based flow cytometry using cells expressing empty vector, full length and Δex5Δex19 NRCAM. Affinity purified IgG2b kappa chains from the hybridoma was further characterized by Western blot analysis and flow cytometry staining.

### Generation of Primary T Cells with FcIR-28z

All primary T cell studies were conducted under approval from the University of Pennsylvania Institutional Review Board (IRB). Identified donors provided informed consent, signing forms approved by the IRB. The activation, transduction, and expansion of primary T cells were carried out using standard techniques. Briefly, primary human CD4+ and CD8+ T cells, obtained from healthy donors through the Human Immunology Core (University of Pennsylvania), were combined in a 1:1 ratio in complete medium (CM) supplemented with 100 IU/mL IL-2 (Prometheus Therapeutics and Diagnostics). On Day 0, the T cells were stimulated with anti-CD3/CD28 Dynabeads (Invitrogen) at a 1:1 bead-to-cell ratio. After 24 hours (Day 1), T cells were transduced with FcIR-28z lentivirus at a multiplicity of infection of 5. The culture volume was doubled daily by the addition of fresh CM until Day 6, at which point the Dynabeads were removed via magnetic separation. Cells were maintained at a concentration of 0.75 × 10□ cells/mL in IL-2 supplemented CM until Day 10, when IL-2 was withdrawn from the medium to complete the expansion process. The efficiency of T cell transduction was assessed on Day 10 by flow cytometry, analyzing for the surface expression of FcIR.

### Cytotoxicity Assays

KNS42 and patient-derived target cell lines were transduced with a pLC-luciferase-zsGreen-blasticidin plasmid and subsequently sorted for GFP-positive populations using fluorescence-activated cell sorting (FACS). Target cells were resuspended in serum-free complete medium (CM) at a concentration of 2x10^5^ viable cells/mL, and 100 µL of cell suspensions was plated per well in a white, opaque-walled, flat-bottom 96-well plate. Cells were cultured overnight at 37°C in 5% CO₂ to reach a confluency of approximately 70-80%. FcIR-28z effector cells were admixed with the antibody 3F8 or its IgG2b isotype control at concentrations of 0, 0.001, 0.01, 0.1, and 1 µg/mL. The “painting” was performed at 37°C in 5% CO₂ for 45 minutes in PBS. Following incubation, effector cells were washed three times in PBS and resuspended in serum-free CM.

To calculate the effector-to-target (E:T) ratio, three random target cell samples from each cell line were trypsinized and counted for live cells. Based on the counts, effector cells were added to the target cells at 3:1, 10:1, and 30:1 E:T ratios. The appropriate number of effector cells was resuspended in 100 µL of serum-free CM and added to the corresponding target cell groups. All conditions were tested in triplicate for each target cell line. Plates were centrifuged at 200g for 3 minutes and then incubated in a humidified incubator at 37°C with 5% CO₂ for 24 hours. Following incubation, 100 µL of medium was carefully removed from each well for further analysis.

For luminescence-based viability assessment, D-luciferin was resuspended in PBS at 900 µg/mL, and 20 µL was added to each well to achieve a final concentration of 150 µg/mL. The plates were placed on an orbital shaker (GeneMate) at 500 rpm for 3 minutes in the dark, then incubated for an additional 3 minutes at 37°C and 5% CO₂. Luminescence readings were obtained using a microplate reader (BioTek Synergy H4), and cytotoxicity was calculated using the following formula: (1 - luminescence (effector + target cells)/luminescence(target cells)) x 100.

## QUANTIFICATION AND STATISTICAL ANALYSIS

### RNA seq analyses

rMATS Turbo (v4.1.0) with GENCODE v39 GFF annotations was used to detect alternative splicing events. Paired comparisons were performed between each tumor sample and seven distinct control sets, including brain homogenate (Clontech, #636643), brain stem (Agilent, #540053, adult), fetal brain (Clontech, #636526), cerebellum (Asterand, #63559-1156128F), and occipital cortex (Asterand, #38061-113046A4). Additionally, publicly available datasets were incorporated, including fetal brain (SRR4787052) and pediatric normal cortex (SRR7268790, SRR7268791, SRR7268792). Aberrant splicing events were filtered to retain those with ≥10 junction read counts. The filtered results served as the basis for all downstream analyses.

For target discovery, we extracted all single-exon (SE) splicing events with a ΔPSI ≥ |0.30| or a ΔPSI ≥ 0.10 when the control PSI was <.10. Additionally, they had to meet significance thresholds (FDR and p-value of < 0.05) in all seven comparisons per tumor sample. We then generated BED files representing exons of interest and intersected them with UniProt protein domain topologies using Bedtools (v2.30.0), which allowed identification of exons corresponding to extracellular domains. Box plots were used to visualize splicing events altered in at least 40% of tumor samples, stratified by exon length into the following categories: 0–30 nt, 31–50 nt, 51–100 nt, and >100 nt. The plots were color-coded to denote splicing preference (i.e., inclusion or skipping). Scatter plots were generated using the ggscatter function to highlight microexons (<50 nucleotides). All downstream analyses were conducted using R v4.4.0, with plots generated using ggplot2 v3.5.1.

For the analysis of isoform expression levels, FASTQ files corresponding to GTEx samples and brain cell subpopulations were downloaded, respectively, from dbGaP (accession number phs000424.v10.p2.c1) and GEO (Project GSE73721) using sratoolkit. Reads were aligned using 2-pass mapping to hg38 genome and GENCODE v39 version. Junctions per million (JPM) were calculated by collating “uniquely mapped reads” column from the “SJ.out.tab” files from the STAR aligner output. Uniquely mapped reads were divided by the total number of reads per sample to normalize these splicing junctions for each sample.

### Orthotopic transplantations

In vivo data in Figure 4H represents a single study. However, considerable effort was expanded to minimize bias, with each replicate treated as a separate experiment. Specifically, each batch of pHGG cells (Empty vector, Full-length NRCAM, NRCAM Δex5Δex19) were harvested from 8 cell culture plates (3x8). We then injected 8 mice per group. Both surgeries and orthotopic injections (24 in total) were performed not serially, but independently throughout the day.

