## Supplemental Figures S1-S6 for "NRCAM variant defined by microexon skipping is a targetable cell surface proteoform in high-grade gliomas"

**Supplemental Figures S1-S6 for D-25-00859R3 "NRCAM variant defined by microexon skipping…" (Sehgal et al)**

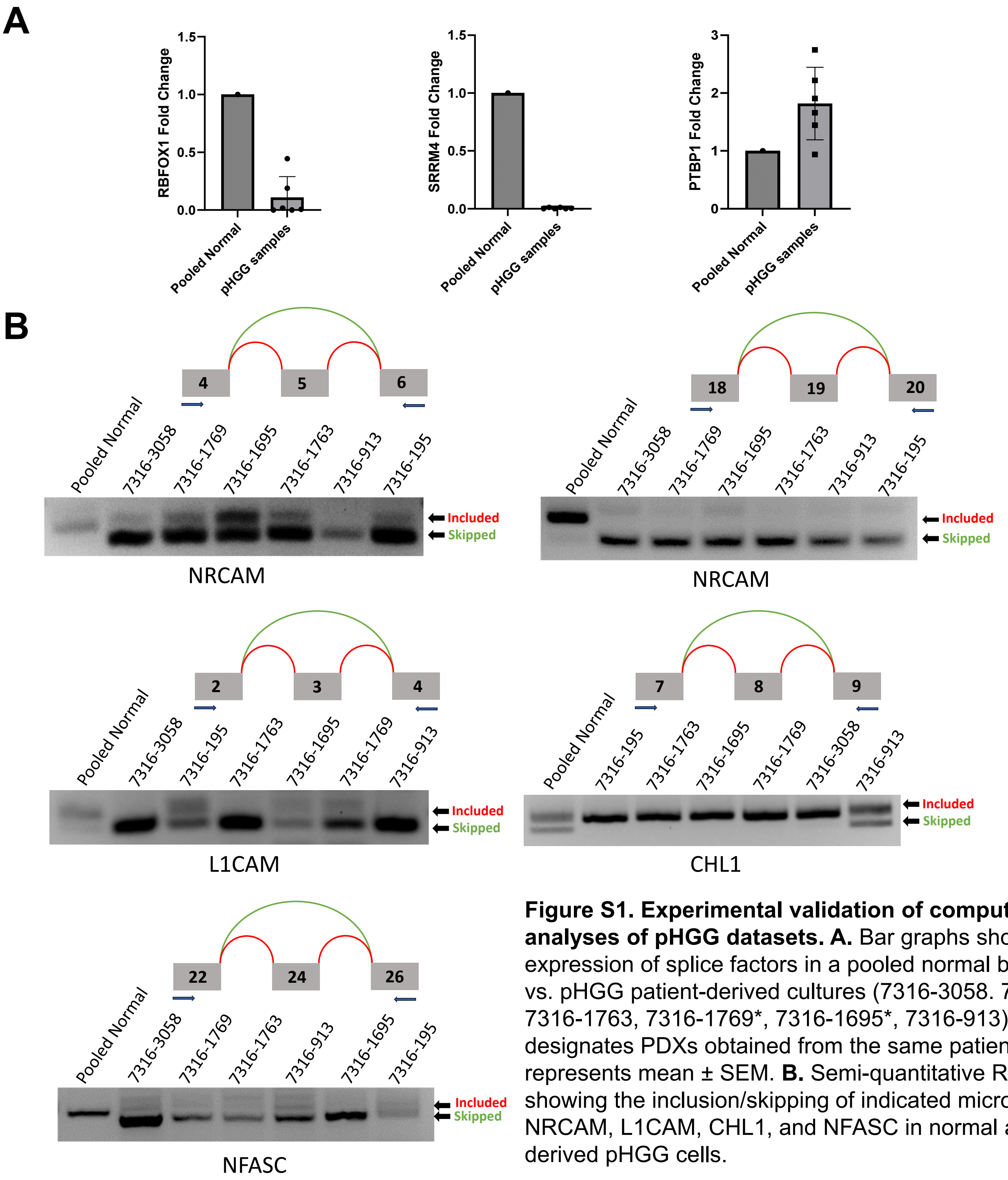


**Figure S1. Experimental validation of computational analyses of pHGG datasets. A.** Bar graphs showing the expression of splice factors in a pooled normal brain sample vs. pHGG patient-derived cultures (7316-3058. 7316-195, 7316-1763, 7316-1769*, 7316-1695*, 7316-913). * designates PDXs obtained from the same patient. Each bar represents mean ± SEM. **B.** Semi-quantitative RT-PCR showing the inclusion/skipping of indicated micro-exons in NRCAM, L1CAM, CHL1, and NFASC in normal and patient-derived pHGG cells.

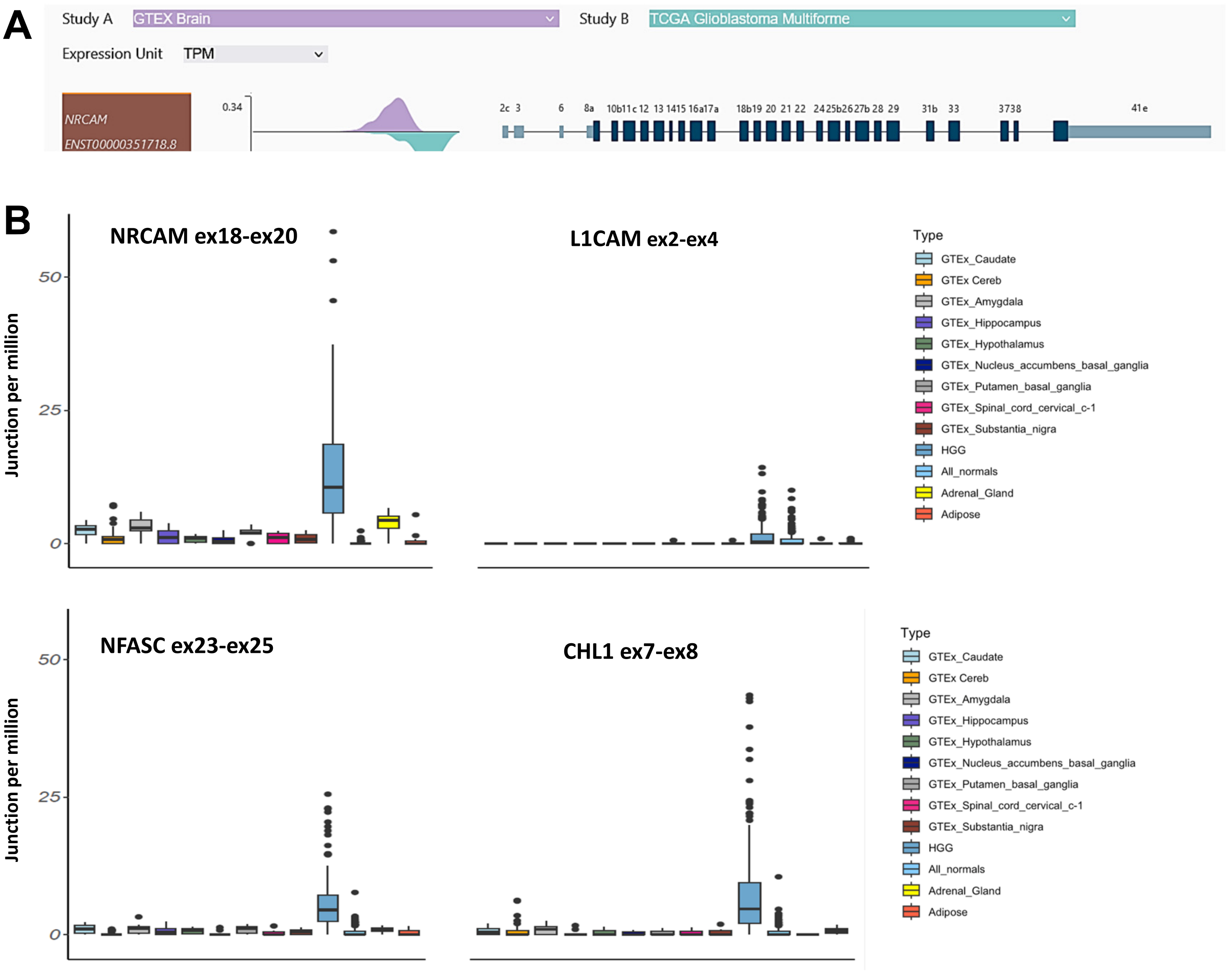


**Figure S2. Expression patterns of L1-IgCAM family members. A.** Visualization of the ENST00000351718.8 transcript and its expression level in normal brain (purple) vs. glioblastoma multiforme (teal) using Xena portal. **B.** Expression of the Δex19 variant of NRCAM mRNA, the Δex3 variant of L1CAM mRNA, the Δex24 variant of NFASC mRNA and the +ex8 variant of CHL1 mRNA in indicated GTEx tissues (neural- and non-neural) and pHGG samples. Read counts corresponding to exon-exon junction were used to estimate transcript levels and are plotted as junctions per million on the *y* axis. Horizontal lines correspond to median values.

**A**

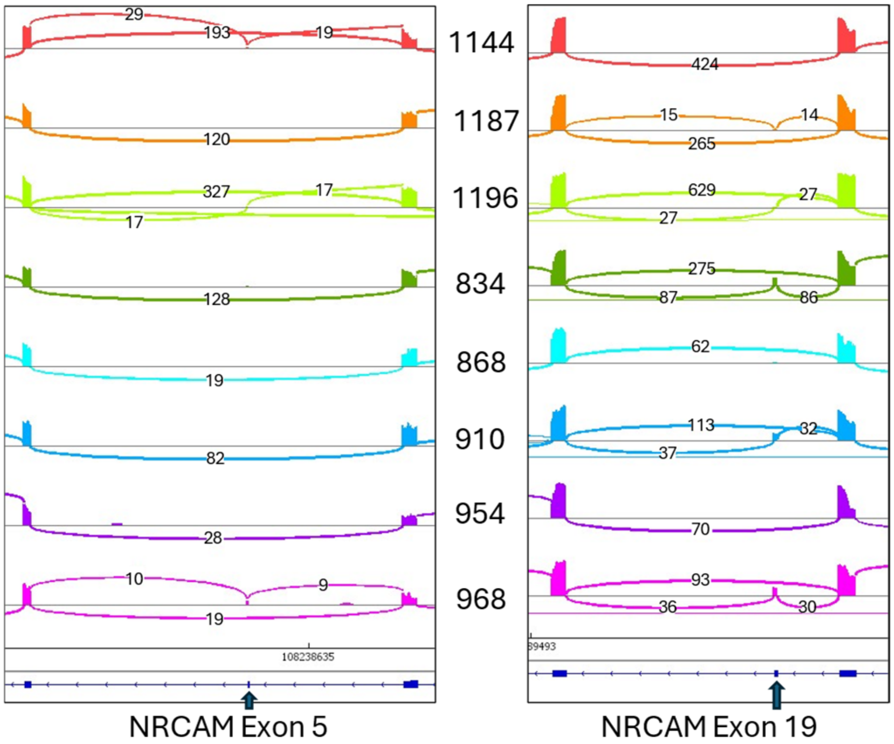


**B**

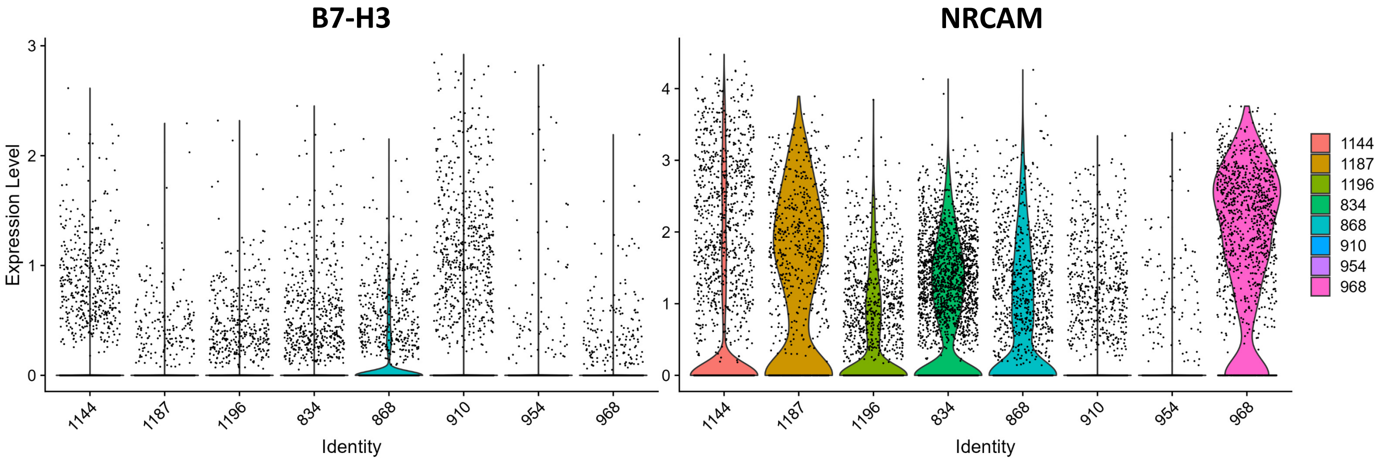


**Figure S3. Bulk and single-cell RNA-seq analysis of 8 pHGG samples from the GSE231859 dataset. A.** Reads corresponding to skipping/inclusion of NRCAM exon 5 and exon 19, visualized in IGV and shown as sashimi plots. **B.** Violin plots showing expression of B7H3 (left) and NRCAM (right) mRNAs across all cells in each sample.

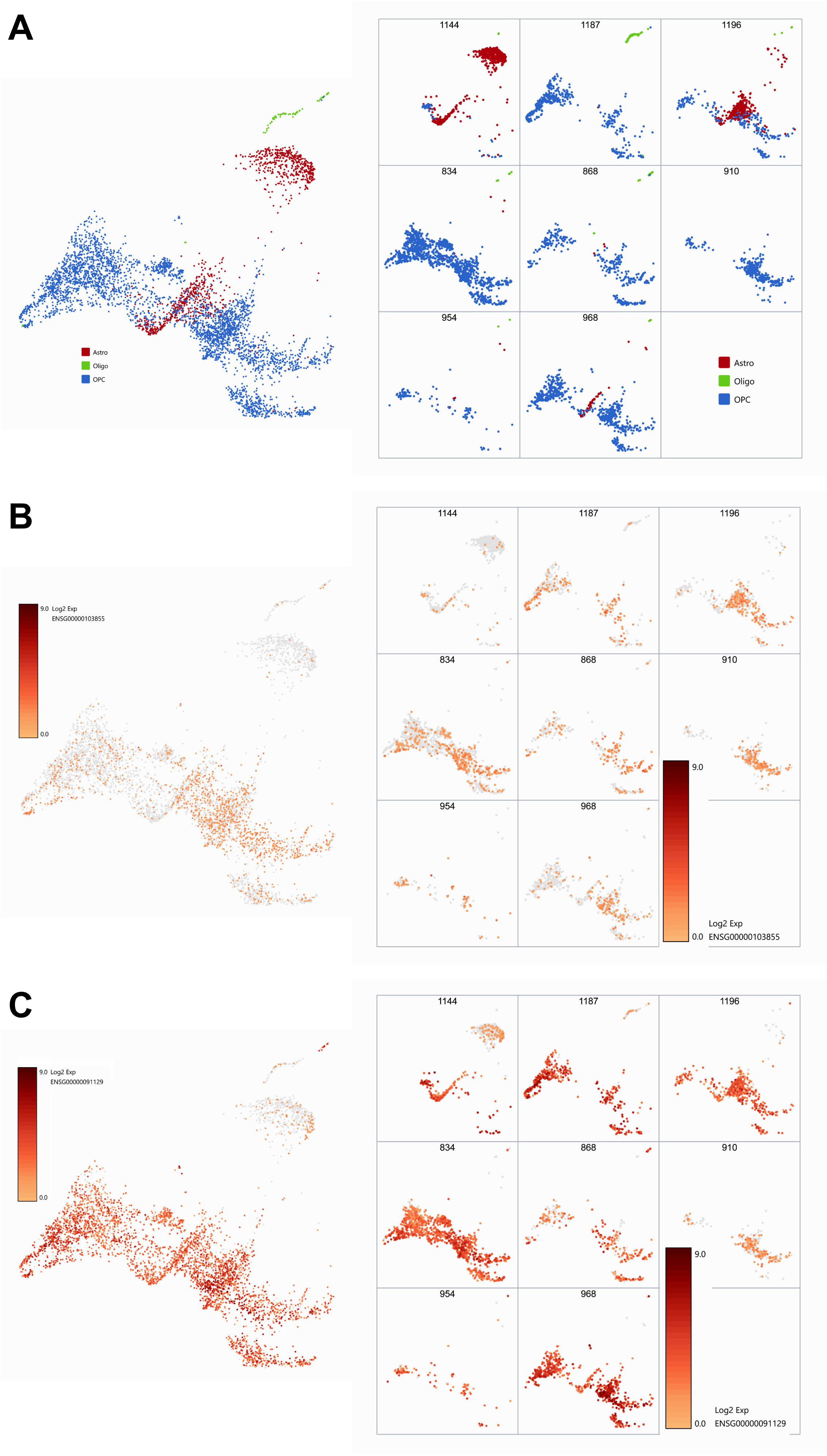


**Figure S4. Single-cell analysis of NRCAM and B7H3 expression in neoplastic cells from the GSE231859 dataset. A.** UMAP projections of pooled sample cells annotated as being of glial origin (astrocytes, oligo-dendrocytes, and OPCs). **B.** UMAP plots of the same cells showing expression levels of B7H3 mRNA. **C.** UMAP plots of the same cells showing expression levels of NRCAM mRNA.

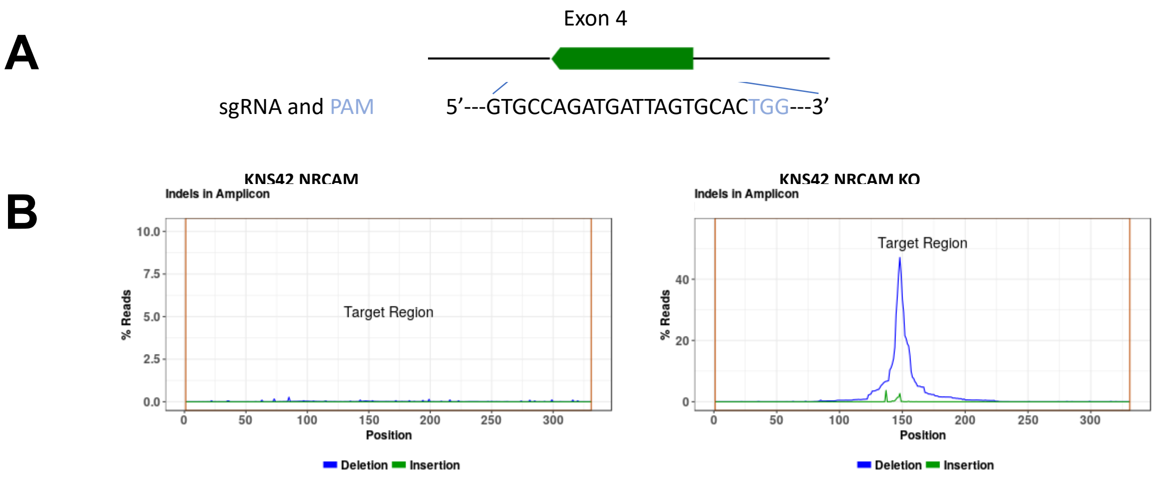


**Figure S5. Generation and validation of NRCAM KO cells. A.** Short guide RNA aligned to NRCAM exon 4 and used to generate Cas9 RNP particles. **B**. Percentage of mutations in parental KNS42 cells vs. cells treated with the sgRNA/Cas9 RNP, as determined by amplicon re-sequencing.

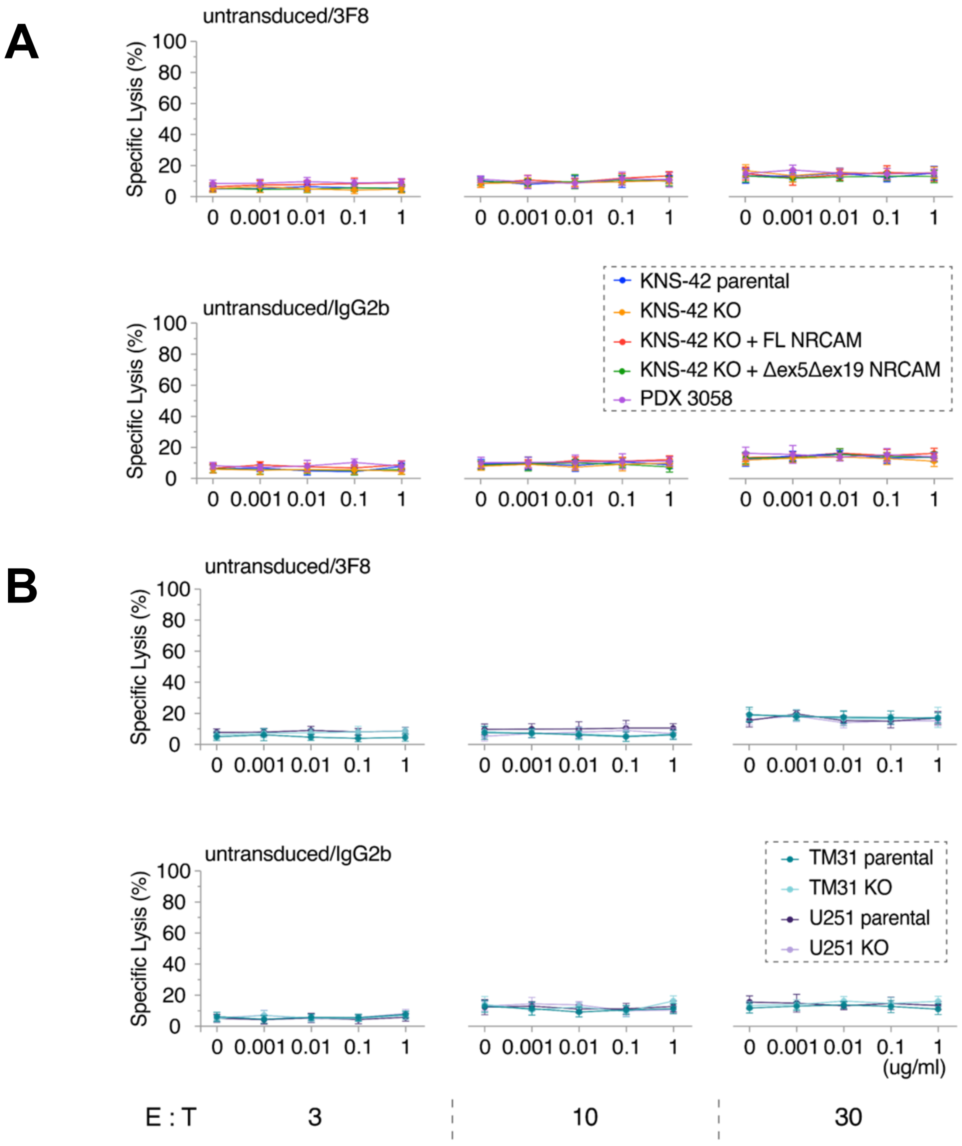


**Figure S6. A. Survival of pHGG and GMB cells treated with untransduced donor T cells. A.** Survival of PDX3058 and KNS42 cells expressing indicated NRCAM isoforms. **B.** Survival of adult glioblastoma U251 and TM31 cells and their NRCAM KO derivatives. In both panels, shown on the *x* axis are Ab concentrations (in mg/ml), and on the *y* axis - the extent of tumor cell killing, as evidenced by reduced luciferase expression. "untr" denoted untransduced T cells, "3F8" – the 3F8 mAb, "IgG2b" – the isotype control. "E:T" values refer to the ratio of effector (T) to target (glioma) cells.
